# Vertical inheritance and loss-driven evolution of secretion systems in the bee gut microbiota

**DOI:** 10.64898/2026.03.23.713742

**Authors:** Samuel A. Acheampong, Waldan K. Kwong

## Abstract

The stability of gut bacterial communities is determined by complex inter-cellular interactions such as competition, cooperation and host dynamics. A mechanism proposed to mediate these interactions is bacterial secretion systems: specialized protein complexes that secrete effector molecules into neighbouring cells or the surrounding environment to influence community stability. However, the forces driving secretion system distribution and evolution in host-associated microbiomes remain unclear. Here, we show that secretion systems in the bee gut microbiome are predominantly vertically inherited and evolve primarily through recurrent gene loss rather than horizontal acquisition. Using comparative genomic analysis, we found that bee gut symbionts mostly encode type I, V, and VI secretion systems. In contrast, the pathogen-associated type II and III systems are missing, but they retain evolutionarily related pili and flagella. We found weak association between the presence of specific secretion systems and the bee hosts, suggesting that these systems are maintained for interbacterial interactions rather than host-specific adaptation. Co-phylogenetic analyses show congruence between bacterial strain phylogenies and most secretion system phylogenies, indicating a vertical mode of transmission. Only a subtype of the type VI system in the *Orbaceae* and the type IV system in *Snodgrassella* spp. show evidence of horizontal transmission. The lack of horizontal transfers means that losses of secretion systems is a permanent evolutionary event in almost all lineages of the bee gut microbiota. Our study provides a uniquely comprehensive analysis of secretion systems across an entire gut bacterial community, giving insight into how microbiomes evolve and maintain functional interactions within host-associated environments.

## Introduction

Many animals and plants harbour host-specific microbiomes composed of bacterial communities that influence host physiology, development and ecological adaptation (McFall-Ngai et al. 2013; Christian et al. 2015; Hou et al. 2022). Within these host-associated environments, bacteria interact continuously with the host and with co-occurring microorganisms to persist. The traits that mediate such interactions, ranging from metabolic exchange to cooperation and competition, are therefore subject to long-term evolutionary pressures shaped by transmission mode, ecological stability, and opportunities for genetic exchange (Ghoul and Mitri 2016; Baishya and Wakeman 2019; Gorter et al. 2020). Yet, how these pressures influence the evolution and persistence of microbial interaction traits in stable host-associated microbiomes remains poorly understood.

One of the traits that mediate these intercellular interactions is the bacterial secretion systems. These are specialized protein complexes that transport a wide range of effector molecules (mainly proteins) across the cell envelope into neighbouring cells or the extracellular environment, facilitating functions such as host colonization, nutrient acquisition, competition & cooperation, and DNA uptake & competence (Aoki et al. 2010; Leo et al. 2010; Zhang et al. 2011; Green and Mecsas 2016; Sassone-Corsi et al. 2016; Harms et al. 2017; Lin et al. 2017; Speare et al. 2018; Luo et al. 2025; Waksman 2025). In Gram-negative bacteria, secretion systems comprise a diverse set of architectures and functions (T1SS–T9SS, Fig. 1A), and have evolutionarily related extracellular appendages such as type IV and tight adherence pili, and flagella (Büttner and He 2009; Green and Mecsas 2016; Cianciotto and White 2017; Lasica et al. 2017; Mignolet et al. 2018; Coulthurst 2019; Li et al. 2019; Meuskens et al. 2019; Spitz et al. 2019; Whitfield and Brun 2024). Although these systems and appendages are considered important determinants of microbial ecology, they have been predominantly studied in pathogenic and free-living bacteria, mostly in isolation or in simplified experimental systems (Green and Mecsas 2016; Nazir et al. 2017; Nicholson and Champion 2022; Maphosa et al. 2023; Jamali et al. 2025). They are often considered highly dynamic traits, frequently shaped by horizontal gene transfer, gene loss, duplications and rearrangements (Abby and Rocha 2012; Denise et al. 2020).

**Figure 1.**
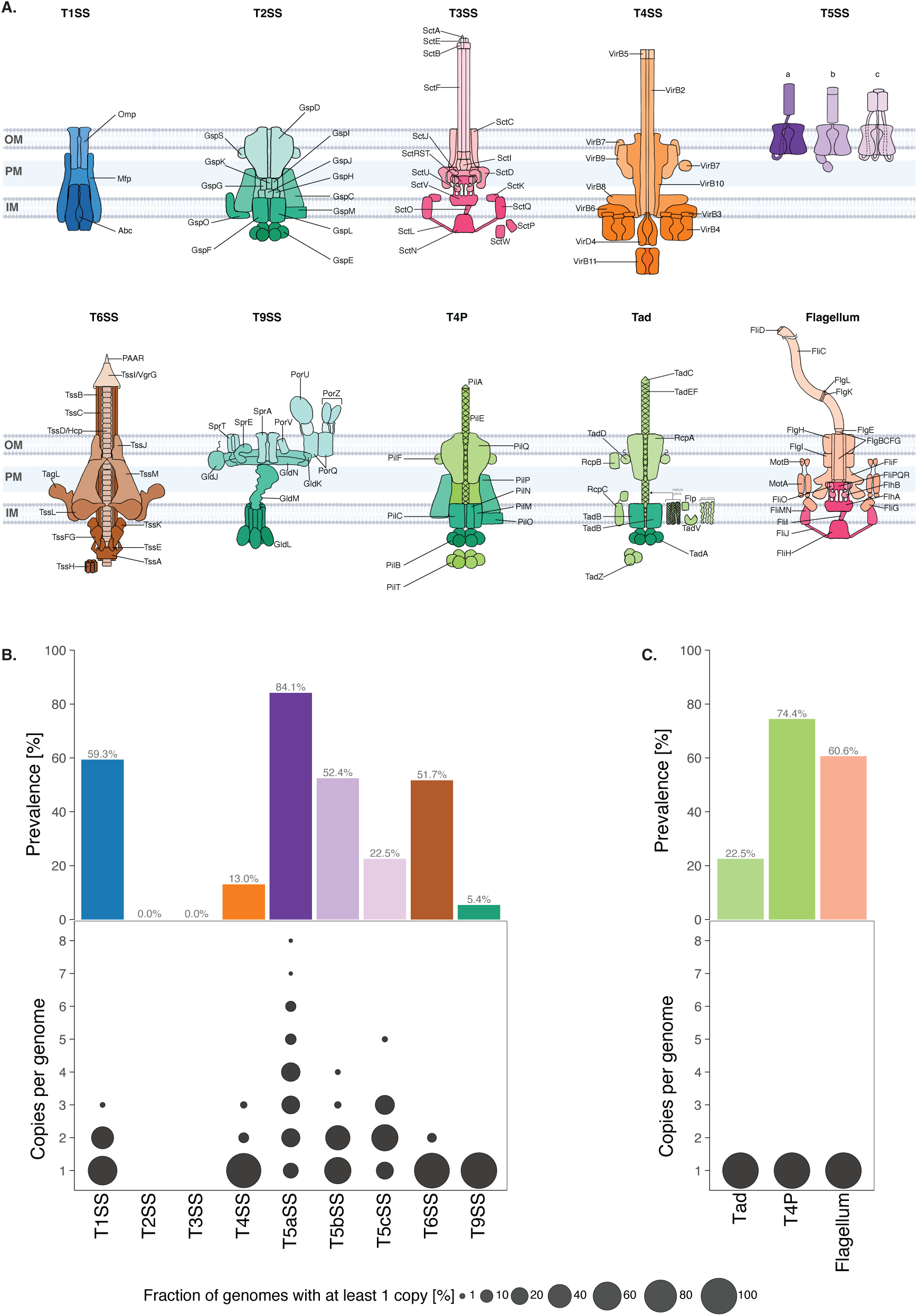
Prevalence of secretion systems and related appendages in bee-associated bacteria. **(A)** Illustration of secretion systems types I (T1SS) to IX (T9SS) and their evolutionary-related appendages type IV pili (T4P), tight-adherence pili (Tad) and flagella, showing putative assembly of individual structural components across the inner membrane (IM), periplasm (PM) and the outer membrane (OM), of Gram-negative bacteria. **(B)** Prevalence (presence or absence) and copy numbers of the secretion systems T1SS – T9SS and **(C)** their evolutionary related appendages among the 391 bee-associated Gram-negative bacterial genomes analysed.

Growing evidence suggests that these secretion machineries are also important in non-pathogenic host associations (even commonly retained in genomically-reduced endosymbionts) (Okazaki et al. 2013; Steele et al. 2017; Speare et al. 2018; Schepers et al. 2021; Suria et al. 2022; Zboralski et al. 2022; Wangthaisong et al. 2023; Bontemps et al. 2024; Motta et al. 2024), a view once encapsulated by the concept of symbionts as “intelligent pathogens” (Deakin and Broughton 2009). Contrary to pathogens and free-living bacteria, bacteria belonging to long-term, host-restricted microbiomes may experience different evolutionary trajectories. These symbiotic environments are often characterized by ecological stability, repeated transmission through the host, and reduced exposure to external microbial gene pools – conditions that tend to limit horizontal gene transfer and promote genome streamlining (Moran et al. 2008; Ganesan et al. 2022; Perreau and Moran 2022). Under such constraints, complex and energetically costly traits such as secretion systems may be selectively retained, modified, or lost depending on their long-term utility within the community. As such, secretion systems can be viewed as ecological traits whose evolutionary trajectory reflects a bacterium’s interactions with its host and co-occurring microorganisms. The question remains whether, in such stable microbiomes, secretion systems are primarily acquired horizontally or inherited vertically and what the relative contributions of gene gains and losses are.

The gut microbiome of eusocial bees, including honey bees, bumble bees, and stingless bees, offers a powerful model system for studying these dynamics. Despite its taxonomic simplicity, the bee gut microbiome is remarkably stable across individual bees, colonies, and generations reflecting strong host–microbe associations and potential long-term coevolution (Kwong and Moran 2016a; Kwong, Medina, et al. 2017; Motta and Moran 2024). The bee gut microbiota supports multiple host functions, including nutrient metabolism, protection against pathogens, and modulation of host immunity (Zheng et al. 2016; Kešnerová et al. 2017; Zheng et al. 2017; Lee et al. 2018; Raymann and Moran 2018; Zheng et al. 2019; Steele et al. 2021; Z. Zhang et al. 2022). Transmission of the microbiome occurs largely through interactions among nestmates, reinforcing long-term host association and limiting exposure to external microbial reservoirs (Powell et al. 2014; Kwong and Moran 2016a). These features make the bee gut microbiome particularly well suited for testing hypotheses about the evolutionary stability, transmission mode, and diversification of bacterial interaction traits.

Previous work on the bee gut microbiome has focused primarily on taxonomic composition, metabolic capabilities, and contributions to host physiology, including nutrition, immune modulation, and protection against pathogens. However, the cellular mechanisms that mediate interactions among symbionts and the evolutionary processes shaping these mechanisms remain comparatively underexplored. While individual secretion systems such as the type VI secretion system (T6SS) have been examined in one or few lineages (Steele et al. 2017; Steele and Moran 2021; Schmidt et al. 2023; Motta et al. 2024; Jones et al. 2025), no systematic comparative analysis has addressed the secretion systems repertoires across the microbiome, how they are transmitted and maintained across the bacterial lineages, and what ecological functions they may serve within the community.

Here, we investigate the mode of transmission and evolution of bacterial secretion systems in the bee gut microbiome using comparative genomics and cophylogenetic analyses of 391 bee-associated bacterial genomes spanning the dominant Gram-negative gut lineages. We characterize secretion system repertoires, assess their phylogenetic congruence, and reconstruct patterns of gain and loss over evolutionary timescales. By integrating metagenomic datasets from bees of different host species and age groups, we further evaluate their distribution and abundance in natural communities. Together, our results reveal an evolutionary trajectory in which secretion systems are predominantly vertically inherited and driven by gene loss rather than horizontal acquisition, providing insight into how stable microbial communities evolve and maintain functional interactions within host-associated environments.

## Results

### Bee gut symbionts encode a restricted secretion system repertoire

To systematically identify and quantify secretion systems in bee-associated bacteria, we developed a customised detection pipeline integrating established computational tools TXSScan and SecReT6 (J. Li et al. 2015; Abby et al. 2016; J. Zhang et al. 2022), with system-specific thresholds calibrated to minimise false positives (Materials and Methods). This approach, coupled with manual curation, allowed for greater sensitivity compared to prior gene survey studies that relied on general-purpose annotation tools such as the NCBI Prokaryotic Genome Annotation Pipeline (PGAP) (Barret et al. 2013; Ugarte-Ruiz et al. 2015; Wallner et al. 2021; Liyanapathiranage et al. 2022; Robinson et al. 2022; Robinson et al. 2023). We analysed 391 high-quality, publicly available genomes from bee gut symbionts and bee-associated environments (e.g. honey- and pollen-derived isolates), screening for the presence and absence of conserved core components of bacterial secretion systems and related appendages (Fig. S1-S5). Phylogenetically related bacteria from non-bee hosts and environmental sources were included as outgroups to enable comparative genomic analyses. Subsequent analyses focused on strains from the five dominant Gram-negative lineages of the bee gut microbiota: *Apibacter* (*Weeksellaceae*), *Bartonella* (*Bartonellaceae*), *Bombella* and *Commensalibacter* (*Acetobacteraceae*), *Frischella* and *Gilliamella* (*Orbaceae*), and *Snodgrassella* (*Neisseriaceae*). No secretion systems were detected in the Gram-positive core members (*Lactobacillus*, *Bombilactobacillus*, and *Bifidobacterium*), and these genomes were therefore excluded from further analyses.

Across the analysed genomes, the type V secretion system (T5SS) was the most prevalent, with 84.1% (329 of 391) encoding the T5aSS (classical autotransporters) subtype, 52.4% the T5bSS (two-partner secretion) subtype, and 22.5% encoding the T5cSS (trimeric autotransporter adhesins) subtype (Fig. 1B). The type I secretion system (T1SS) was the second most common, present in 59.3% of the genomes, while the type VI secretion system (T6SS) was found in 51.7%. The T9SS was the rarest, found in only 5.4% of the genomes. To assess copy number variation within genomes, candidate secretion system loci were evaluated using a completion-based criterion requiring the presence of mandatory core components, while accounting for both single-locus and multi-locus architectures defined in the system models (Materials and Methods). Most genomes encoding a given secretion system carried a single copy (Fig. 1B; Fig. S6). In contrast, T5aSS, encoded by a single gene, frequently occurred in multiple copies, with up to eight copies per genome. This pattern was not lineage-restricted, as multi-copy T5aSS were detected across multiple bee-associated lineages (Fig. S6). By contrast, *Bartonellaceae* and most *Acetobacteraceae* genomes typically encoded two copies of the T1SS, whereas *Orbaceae* and *Neisseriaceae* predominantly carried a single copy. Multi-copy T5cSS was restricted to *Neisseriaceae*, which also consistently encoded multiple T5aSS genes (Fig. S6).

### Virulence-associated secretion systems are absent, but related appendages are widespread

The type II secretion system (T2SS) and type III secretion system (T3SS) are commonly associated with pathogenicity (Cianciotto and White 2017; Maphosa et al. 2023). They are known to inject virulence factors into hosts cells, enabling tissue invasion. Strikingly, none of the bee gut or bee-associated bacteria encoded either system (Fig. 1A), suggesting the bee gut and bee-associated bacteria may be lacking host tissue invasive traits. Instead, we observed the widespread presence of the T2SS-evolutionarily related appendages, tight adherence (Tad) pili and type IV pili (T4P) (Peabody et al. 2003) in 22.5% and 74.4% of genomes, respectively and T3SS-related flagellar apparatus (Grognot and Taute 2021; Akahoshi and Bevins 2022) was encoded by 60.6% of genomes (Fig. 1B). These appendages have been implicated in mediating biofilm formation, host attachment, surface sensing, surface motility and inter-bacterial aggregation (Mignolet et al. 2018; Ligthart et al. 2020; Grognot and Taute 2021; Akahoshi and Bevins 2022; Geiger et al. 2024; Whitfield and Brun 2024). All genomes encoding Tad, T4P, or flagella carried a single copy of these appendages (Fig. 1B; Fig. S6).

### Secretion system repertoires are determined by bacterial lineage rather than host identity

Given that secretion systems and appendages often reflect niche adaptation and are non-randomly distributed across bacterial species (Green and Mecsas 2016), we examined their occurrence across the genomes to identify potential host- or lineage-specific patterns in system repertoires. We observed marked lineage-specificity in the distribution of many systems (Fig. 2A). T1SS was highly prevalent in all genera except *Apibacter*. The T2SS-related Tad is present only in the *Orbaceae* family (*Frischella* and *Gilliamella*), while the T4P is present only in *Frischella*, *Gilliamella*, and *Snodgrassella*. T5cSS was found mostly in *Snodgrassella* and T9SS was only carried by *Apibacter*, suggesting phylogenetically constrained functions. Overall, *Gilliamella* and *Snodgrassella* carried the most diverse and variable secretion system repertoires, while *Apibacter*, *Bartonella*, *Bombella*, and *Commensalibacter* possessed highly conserved and streamlined profiles.

**Figure 2.**
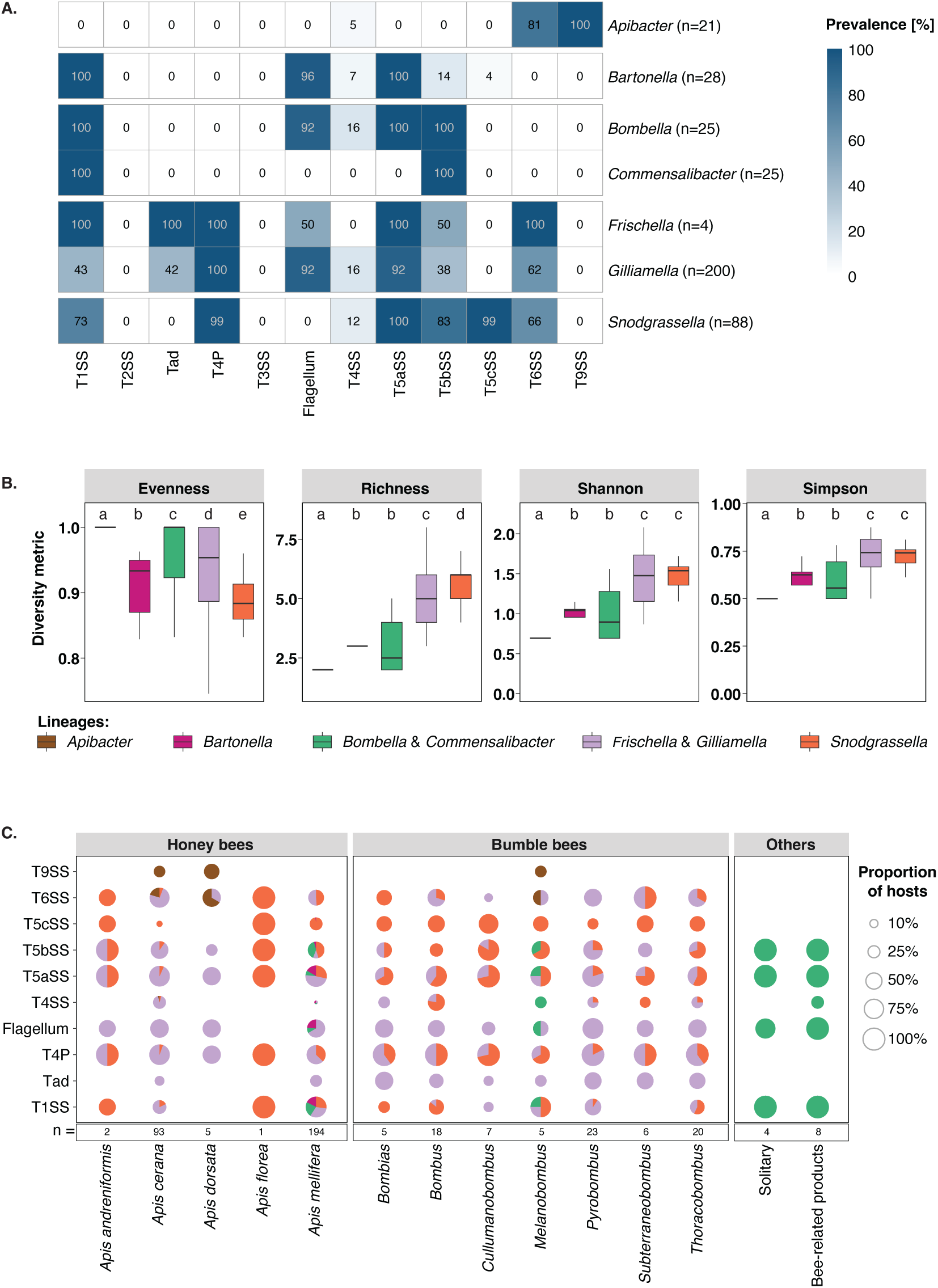
Secretions system occurrence across bacterial lineages and bee hosts. **(A)** Prevalence, given as percentages, of secretion systems and related appendages among each of the Gram-negative bee bacterial genera. Number of strains analysed indicated in parentheses. **(B)** Diversity of secretion system repertoires within each lineage, as assessed with alpha diversity metrics. Pairwise significance determined with Wilcoxon rank-sum tests with Benjamini-Hochberg adjustment; letters indicate groups not significantly different. **(C)** Prevalence of secretion systems and related appendages across different bee hosts. Bee species (for honey bees) and subgenera (for bumble bees) categorized in columns, with n indicating the total number of bacteria originating from each host or source. Pie size indicates the fraction of bacteria per host carrying each system, and the slices show the bacterial lineage contribution. Colours of the slices correspond to the legend in (B).

To quantify these differences, we analysed the alpha- and beta-diversity of the secretion system repertoires of the strains (Fig. 2B, Fig. S7). *Frischella*, *Gilliamella*, and *Snodgrassella* exhibited significantly higher richness, Shannon, and Simpson diversity than the other lineages (Kruskal-Wallis, *p* < 0.01), consistent with their broader range of encoded systems. However, evenness values within these lineages remained moderate, indicating that, although they possess a larger repertoire overall, their profiles are skewed toward a core set of dominant systems that occur in most strains, and additional systems are present only sporadically. In contrast, *Apibacter*, *Bartonella*, *Bombella*, and *Commensalibacter* exhibited both low richness and high uniformity, consistent with their streamlined repertoires. PCoA and beta-dispersion analyses using both abundance and presence/absence data showed a strong lineage effect on secretion system composition, with lineage explaining 68.6% of the variance among strains (Fig. S7).

To assess whether secretion systems show non-random associations across strains, we computed pairwise co-occurrence using Pearson correlations on presence–absence matrices (Fig. S8A). Overall, the co-occurrence analysis revealed no strong positive correlations among secretion systems indicating that the presence of one system provides little predictive information about the presence of another. A few patterns were nevertheless detectable: for example, moderate positive associations between Tad, T4P, and the flagellum, and between T4P and T5aSS, whereas T5cSS and the flagellum exhibited a strong negative association. Considering that the dataset is heavily dominated by *Frischella* & *Gilliamella* (n = 204) and *Snodgrassella* (n = 88), and that lineage-dependence (i.e., phylogenetic signal) could influence the co-occurrence patterns, we evaluated associations while explicitly accounting for phylogeny using Coinfinder v1.2.1 (Whelan et al. 2020). This approach estimates whether co-occurrence is more frequent than expected after controlling for shared evolutionary history. In *Frischella* & *Gilliamella*, we found that T1SS, Tad, and T6SS tended to co-occur more often than expected irrespective of phylogeny (*p* < 0.05, Fig. S8B). In *Snodgrassella*, T1SS and T5bSS showed a weak but non-significant pattern of co-occurrence (*p* = 0.08, Fig. S8C).

Since the analysed bacterial strains originate from a variety of bee host species, and strains can be highly specific to their particular bee hosts (Kwong and Moran 2016a; Powell et al. 2016; Kwong, Medina, et al. 2017; Ellegaard et al. 2020), we considered if secretion systems also display host specificity. When mapped to host identity, we found patterns of host-specificity for only a few systems, supported by weak clustering in PCoA analysis (Fig. 2C, Fig. S9). T9SS was present only in *Apis cerana*, *Apis dorsata*, and the bumble bee subgenus *Melanobombus*, with this distribution driven by the specificity of *Apibacter* strains to these hosts. Intriguingly, the T4SS appeared to be enriched in bumble bee derived strains compared to strains from honey bees. The T4SS is infrequently found in bee gut strains overall (Fig. 1B) but is sporadically present in members of all 5 analysed bee gut lineages.

### Secretion systems are predominantly vertically inherited

To assess whether secretion systems were vertically inherited or horizontally acquired, we compared the phylogenies of secretion systems to their corresponding bacterial lineage phylogenomic trees. Phylogenetic congruence, as a proxy of mode of inheritance, was evaluated using three distance-based cophylogenetic methods, Parafit, Hommola and PACo, and one event-based method, eMPRess (Legendre et al. 2002; Hommola et al. 2009; Hutchinson et al. 2017; Santichaivekin et al. 2021). Significant congruence in these phylogenies would suggest co-divergence of secretion systems with the bacterial strains. Across most systems and lineages, we observed significant phylogenetic congruence (*p* < 0.05) in at least two of the distance-based methods and in the event-based method (Table 1), consistent with vertical inheritance. To visually validate these results, we constructed species-system tree tanglegrams and heatmaps of pairwise amino acid identity from the alignments used to build each secretion system tree.

**Table 1.** Co-phylogeny analyses of bacterial lineages and their secretion systems. *P*-values reported from three separate distance-based methods (PACo, ParaFit, Hommola) and one event-based method (eMPRess) to assess congruence between bacterial lineage phylogenies and system phylogenies. Phylogenies were constructed with conserved, single copy orthologs (details in Materials and Methods). Values in bold indicate lack of support for phylogenetic congruence (*p* > 0.05).

| Lineage | System | *p*-value |  |  |  |
| --- | --- | --- | --- | --- | --- |
|  |  | Parafit | Hommola | PACo | eMPress |
| *Apibacter* | T6SS | 0.001 | 0 | 0 | 0.020 |
|  | T9SS | 0.002 | 0 | 0 | 0.010 |
| *Bartonella* | **T1SS (both clades)** | **1.000** | **0.870** | **0.999** | 0.010 |
|  | T1SS clade 1 | 0.001 | 0 | 0 | 0.010 |
|  | T1SS clade 2 | 0.001 | 0 | 0 | 0.010 |
|  | Flagellum | 0.001 | 0 | 0 | 0.010 |
|  | T5aSS | 0.015 | 0.010 | 0.004 | 0.010 |
| *Bombella & Commensalibacter* | T1SS | 0.001 | 0 | 0.001 | 0.010 |
|  | Flagellum | **0.411** | 0 | 0 | 0.010 |
|  | T5aSS | **0.915** | 0 | 0 | 0.010 |
|  | T5bSS | 0.001 | 0 | 0 | 0.010 |
| *Gilliamella & Frischella* | T1SS | 0.001 | 0 | 0 | 0.010 |
|  | T4P | 0.002 | 0 | 0 | 0.010 |
|  | Tad | 0.001 | 0 | 0 | 0.010 |
|  | Flagellum | 0.005 | 0 | 0 | 0.010 |
|  | T4SS | 0.022 | 0 | 0 | 0.011 |
|  | T5aSS | 0.001 | 0 | 0 | 0.010 |
|  | T5bSS | 0.001 | 0 | 0 | 0.010 |
|  | T6SS (both subtypes) | 0.001 | 0 | 0 | 0.010 |
|  | **T6SS subtype i1** | 0.019 | **0.055** | 0.045 | **0.257** |
|  | T6SS subtype i2 | 0.001 | 0 | 0 | 0.010 |
| *Snodgrassella* | T1SS | 0.001 | 0 | 0 | 0.010 |
|  | T4P | 0.001 | 0 | 0 | 0.010 |
|  | **T4SS** | **0.669** | **0.877** | **0.459** | **0.327** |
|  | T5aSS | 0.001 | 0 | 0 | 0.010 |
|  | T5bSS | 0.001 | 0 | 0 | 0.010 |
|  | T6SS (both subtypes) | **0.068** | 0 | 0.007 | 0.010 |
|  | T6SS subtype i2 | 0.001 | 0 | 0 | 0.010 |
|  | T6SS subtype i3 | 0.001 | 0 | 0 | 0.010 |

In most cases, the congruence tests and heatmap visualizations were consistent. The T1SS in *Bartonella* appeared to be an exception, with congruence tests suggesting horizontal transmission. However, upon closer examination, we found that all bee-associated *Bartonella* strains encoded two distinct T1SS copies that formed two well-supported clades, each corresponding to one of the copies. When these clades were analysed separately, both exhibited signatures of vertical inheritance (Table 1, Fig. S10). Phylogenetic topology and additional sequence alignments indicated that these copies are unlikely to represent recent paralogs arising from gene duplication, suggesting instead that they may have independent evolutionary histories. A BLAST search revealed the closest hits of clade 1, at ≥55% amino acid similarity, were from *Bartonella* strains of other hosts such as humans (*Bartonella tamiae*) and bats (*Bartonella sp. HY406*), while that of the clade 2, at ≥30% amino acid similarity, were from strains of distant Beta- and Gammaproteobacteria. This suggests that the clade 1 T1SS was vertically inherited from an ancestral *Bartonellaceae,* while the clade 2 T1SS may have been acquired horizontally by a common ancestor prior to, or during the early stages of association with bees, and transmitted vertically afterwards.

### Horizontal acquisition of T4SS and T6SS in two lineages

Exceptions to the pattern of strict vertical inheritance were observed for the T4SS of *Snodgrassella* and the T6SS subtype i1 within the *Frischella*/*Gilliamella* lineage (Table 1). In *Snodgrassella*, the T4SS phylogeny showed topological incongruence with the species tree (Fig. 3A) and pairwise T4SS nucleotide similarity was lower and far more variable (mean = 70.1% ± 21.41%) than the overall average nucleotide identity (ANI) among the strains (mean = 93.13% ± 6.35%) (Fig. 3B). Notably, *Snodgrassella sp.* B3088 harbours two distinct, highly divergent T4SS loci. BLAST analysis shows the closest hit to locus1 T4SS is the conjugal T4SS from a relative within the *Neisseriaceae*, *Neisseria flavescens* (72.1% identity), while locus2 matches genes from *Paralysiella testudines* (70.8%). This suggests the copies arose from independent acquisition rather than duplication. Furthermore, the two *Snodgrassella communis* strains LMG28360 and R-54236 share 100% T4SS identity to one another but very low identity (<40%) to the T4SS of the other strains, including other *S. communis* strains, suggesting another instance of independent horizontal acquisition.

**Figure 3.**
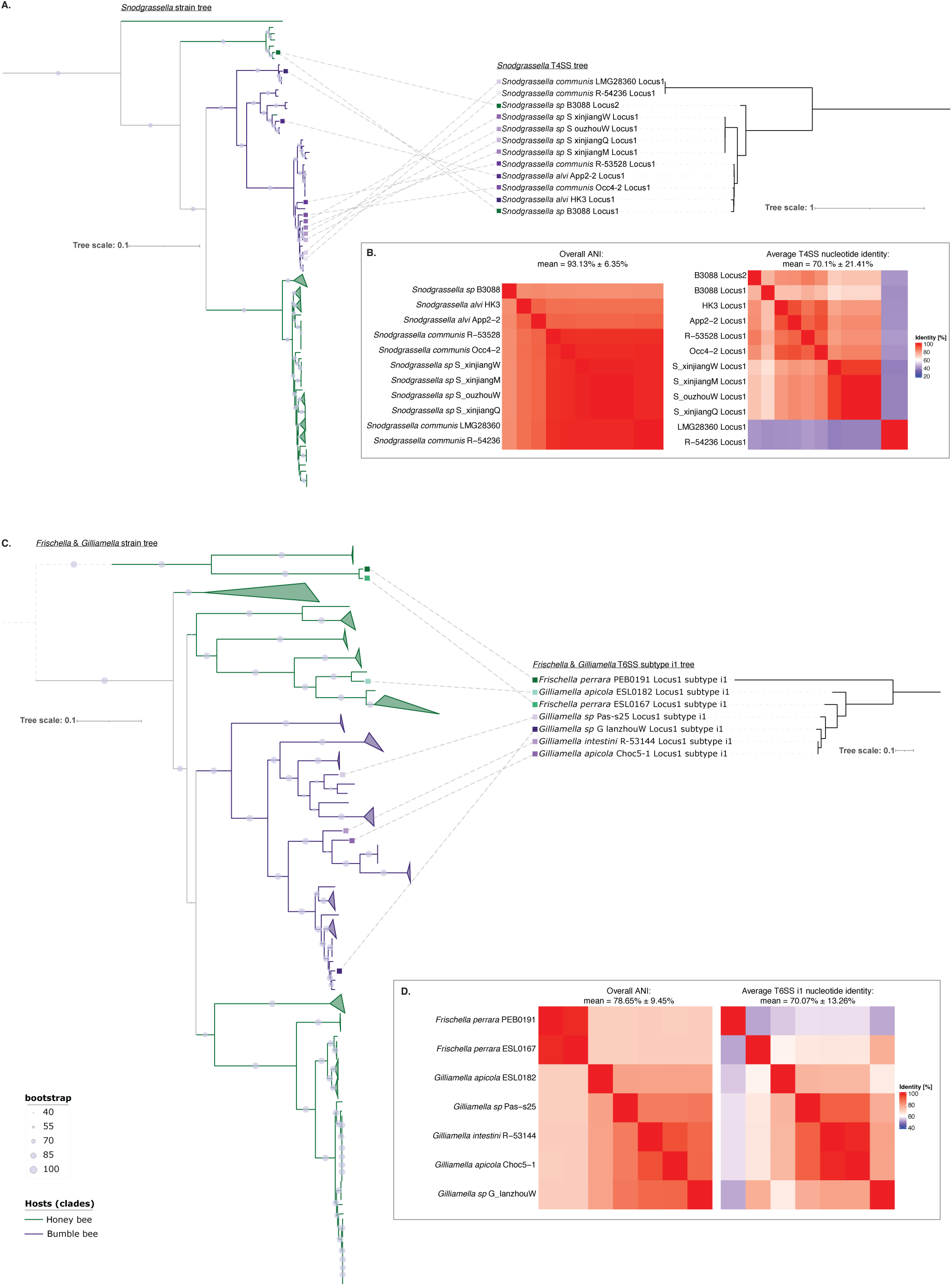
Two exceptions to the vertical inheritance of secretion systems in the bee microbiota. **(A)** Tanglegram between *Snodgrassella* strain phylogeny and the type IV secretion system phylogeny, showing phylogenetic incongruence. **(B)** Pairwise average nucleotide identity (ANI) between *Snodgrassella* genomes and between their T4SS genes. **(C)** Tanglegram between *Frischella* & *Gilliamella* strain phylogeny and the type VI secretion system subtype i1 phylogeny. **(D)** Corresponding genomic ANI and ANI of their T6SS genes. Tree node support values are shown as bootstrap percentages.

Two of our four cophylogenetic tests revealed incongruence specifically in T6SS subtype i1 within the *Frischella*/*Gilliamella* lineage, which is supported by tanglegram visualization (Fig. 3C). The T6SS is categorised into four subtypes (T6SSi-iv) of which the T6SSi is further subdivided into i1, i2, i3, i4a, i4b, and i5 (Boyer et al. 2009). *Frischella* and *Gilliamella* generally encode a single copy of the T6SS, the majority of which (123/204 strains) corresponds to subtype i2, and which appears to be vertically inherited (Table 1). Subtype i1 is restricted to five *Gilliamella* strains, where subtype i1 is present instead of i2, and two *Frischella* strains, which carry both subtypes. While *Frischella* strains PEB0191 and ESL0167 cluster together in the species tree (98.8% ANI), their subtype i1 T6SSs are highly divergent, sharing only 52.22% nucleotide similarity (Fig. 3D). Conversely, *Gilliamella* strains R-53144 and Choc5-1 show the opposite pattern: despite being more divergent at the genome level (94.0% ANI), their T6SS are nearly identical (99.0% nucleotide similarity). This is consistent with horizontal transfer of T6SS between these strains, but while some T6SSs are known to be encoded on mobile genetic elements (García-Bayona et al. 2021; Li et al. 2021; Jana et al. 2022; Unni et al. 2022), we found no clear evidence of transposon or plasmid-related genes near the subtype i1 locus in these bee symbionts. Nonetheless, the phylogenetic incongruence and discordant sequence similarities suggests the T6SS subtype i1 can be transmitted horizontally in *Frischella*/*Gilliamella*.

### Gene loss is the dominant force driving secretion system evolution

To further explore the evolutionary history of these systems, we mapped the binarized distribution of each detected system onto the respective lineage-specific phylogenomic tree and examined patterns of predicted gain and loss events (Fig. 4). Overall, the reconstruction of events shows that secretion system evolution is dominated by repeated losses rather than gains. For instance, although the T1SS is highly prevalent, across both closely and distantly related species within the *Weeksellaceae*, it is entirely absent in the bee *Apibacter* strains (Fig. 4A) suggesting that its loss likely occurred prior to *Apibacter* diversification. In contrast, the T6SS is markedly enriched in bee *Apibacter* strains compared to their relatives. However, within *Apibacter*, several strains lack T6SS, indicating more recent and likely independent loss events.

**Figure 4.**
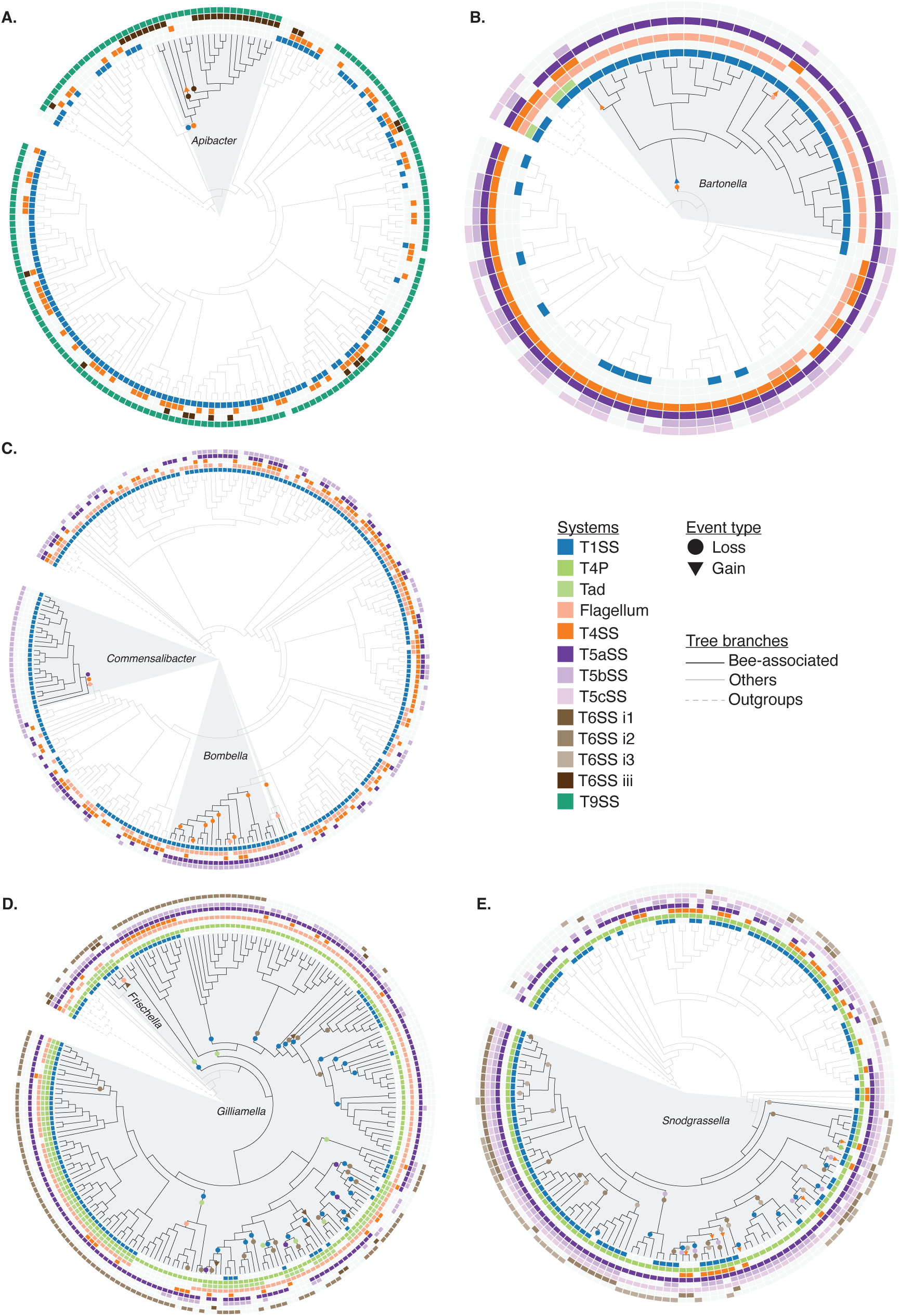
Secretion system losses and gains over the evolution of the bee microbiota. Phylogenetic trees of bee-associated bacterial lineages and related taxa (see Figs. S1–5 for detailed trees) of **(A)** *Apibacter*, **(B)** *Bartonella*, **(C)** *Bombella* and *Commensalibacter*, **(D)** *Frischella* and *Gilliamella*, and **(E)** *Snodgrassella*. The bee bacteria clades are shaded in grey. The presence and absence of each secretion system is shown in tracks around the phylogenetic trees. Loss and gain events are shown as filled shapes on the inferred clades/branches of the event.

Possible rare cases of secretion system gain were observed in the *Bartonellaceae*: two strains possessed the T4SS despite it appearing to have been lost early in the bee *Bartonella* lineage (Fig. 4B). These two T4SSs were most similar to those of *Brucella/Ochrobactrum* and the plant root bacteria *Mesorhizobium*, suggesting that they were recent gain events. For *Bombella* and *Gilliamella*, the scattered distribution of their T4SS (Fig. 4C, D), and the lack of clear comparisons with related taxa makes it difficult to deduce their loss or gain events; nonetheless, their closest BLAST hits were to the conjugal T4SS in bacteria such as *Pseudomonas*, *Frigidibacter*, *Acetobacter* and *Salmonella*, which may indicate horizontal acquisition events early in the evolution of these bee-associated lineages. In *Gilliamella* and *Snodgrassella*, multiple independent loss events of the T1SS (25 events in *Gilliamella*, 10 in *Snodgrassella*), T4P (7 events in *Gilliamella*, 1 in *Snodgrassella*) and T6SS (18 events in *Gilliamella*, 27 in *Snodgrassella*) appear to have occurred, but predominantly after the diversification of the bee lineages. The absence of the T1SS, Tad, and T6SS in several divergent branches suggests recent, independent loss events within these lineages. *Snodgrassella* strains typically encode two T6SS: subtype i3 at single genomic locus and subtype i2 that is often split across two separate loci. Both showed strict vertical inheritance (Table 1), in agreement with a previous analyses (Steele et al. 2017). Our phylogenetic reconstruction indicates that a post-diversification ancestor likely encoded both subtypes (Fig. 4E). Multiple independent gene loss events within the lineage subsequently produced strains harbouring either zero, one, or both subtypes.

Overall, while rare gains do occur via horizontal gene transfer, system losses are far more frequent and typically irreversible.

### Secretion system abundances vary with host age and species

While our genome-based comparative analysis revealed the prevalent secretion systems encoded by the bee gut and bee-associated bacterial genomes, it does not capture their diversity or distribution within natural microbial communities. To address this, we analysed three publicly available metagenomic datasets derived from *Apis cerana* and *Apis mellifera* hosts of varying ages and geographic origins (Ellegaard and Engel 2019; Ellegaard et al. 2020; Wu et al. 2021). We constructed a curated database of secretion system component sequences identified from our genome analysis and used these as queries against the metagenomes under stringent thresholds and thus were able to differentiate between secretion systems encoded by bee specific bacteria and those from other low abundance or transient bacteria. Adjusted for abundance (read depth), most detected secretion systems in metagenomes belonged to members of the bee microbiome (Fig. S11-S13).

Across the metagenomes, the taxonomic distribution of secretion system components from bee bacterial lineages largely mirrored the patterns observed in the genomes (Fig. S14). For example, T9SS was exclusively associated with *Apibacter*, Tad systems with *Frischella* and *Gilliamella*, and T1SS was broadly distributed across most taxa except *Apibacter*. As expected from their dominance among Gram-negative bacteria in individual gut communities, components originating from *Gilliamella* and *Snodgrassella* were highly represented (Fig. S14, S15). Among all metagenomic reads assigned to secretion systems in the dataset from Ellegaard et al. (2019), 60.37% were assigned to *Gilliamella*, 13.04% were assigned to *Snodgrassella*, 21.25% were assigned to *Bartonella*, while fewer than 0.01% were assigned to *Apibacter*, predominantly corresponding to the T9SS. Similar patterns were observed in the datasets from Ellegaard et al. (2020) and Wu et al. (2021), in which 73.06% and 51.68% of the total assigned reads, respectively, corresponded to *Gilliamella*, while 12.19% and 19.93%, respectively, corresponded to *Snodgrassella*.

We next examined whether secretion system diversity varied as a function of host age (Fig. 5). To quantify secretion system abundance, we mapped raw reads to their respective components, normalised by gene length and sequencing depth, and expressed the results as copies per million (CPM). Overall, secretion system diversity declined significantly with host age (Fig. 5A). This pattern was not attributable to shifts in the diversity of contributing taxa, which did not differ significantly across age groups for any classified taxon. Younger bees harboured more even profiles of secretion systems, reflected in higher Shannon and Simpson diversity indices, whereas older bees were dominated by a smaller set of systems, particularly flagella and T5SS (Fig. 5A). To assess the extent to which taxonomic composition contributed to these age-associated differences, we calculated the relative abundance of each secretion system after normalising by the abundance of its contributing taxa (Fig. S16). The decline in T6SS with age occurred mainly within *Snodgrassella* and *Gilliamella*, whereas increases in T1SS, Tad, and flagella were driven largely by *Bartonella*, *Commensalibacter*, and *Frischella*. These patterns suggest that age-related changes in secretion system composition arise not only from shifts in genus-level community structure but also from differences among strains within genera. Functionally, the elevated abundance of Tad, T4P, and T6SS in young bees suggests that early gut communities are primarily engaged in processes of colonization and interbacterial competition. In contrast, the increased abundance of flagellar systems, T1SS, and T5SS in older bees may indicate a functional shift toward motility, sensing and possibly resource exploitation, and host-associated interactions.

**Figure 5.**
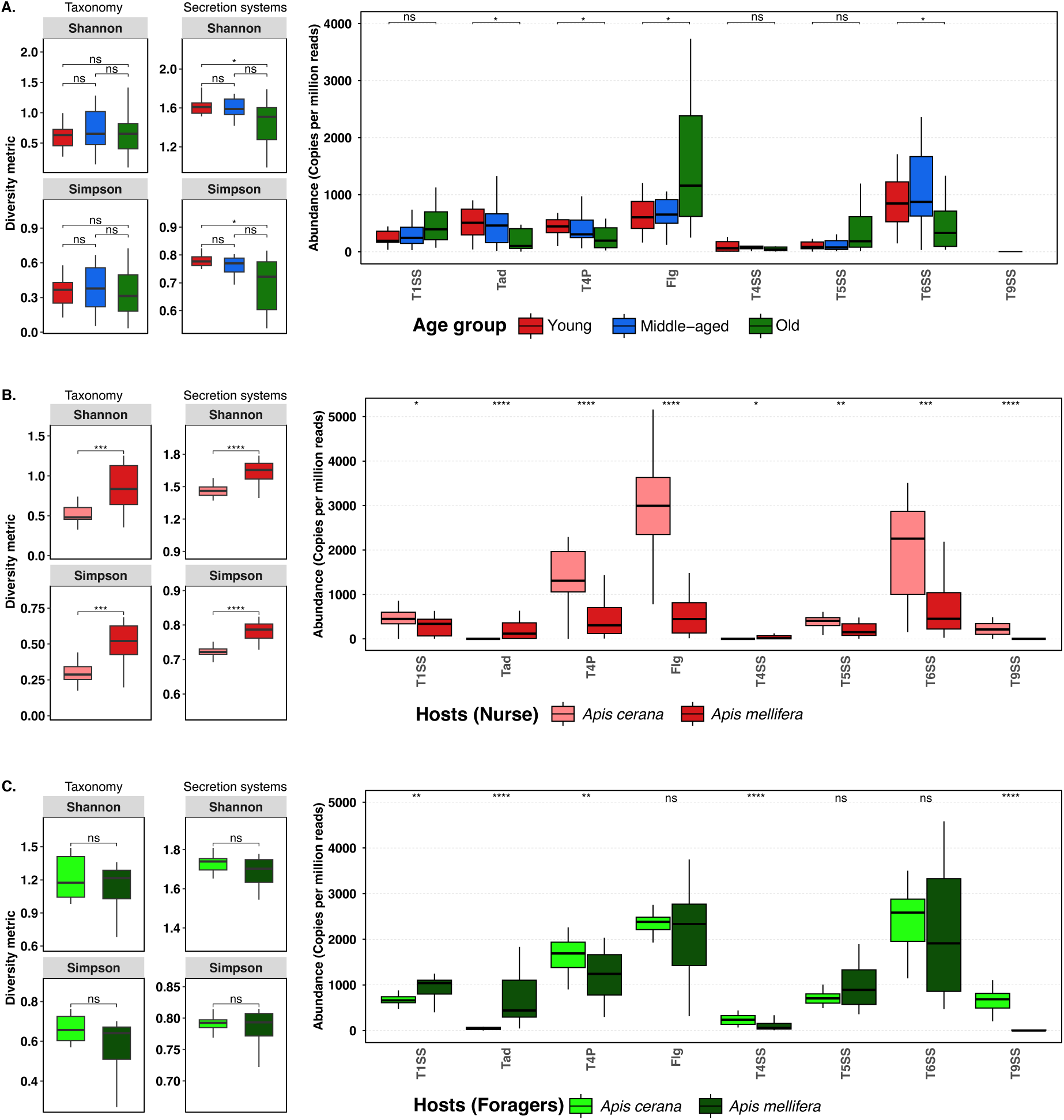
Influence of host age on secretion system diversity. Taxonomic diversity and secretion system diversity of bee-associated Gram-negative bacteria from metagenomic samples, and normalized abundances of reads corresponding to each secretion system within samples. **(A)** Young (10 days old), middle-aged (22–24 days) and old (≥48 days) *A. mellifera* (Ellegaard *et al*., 2019); **(B)** from nurses (young) *A. cerana* and *A. mellifera* (Ellegaard *et al*., 2020); and **(C)** from foraging (old) *A. cerana* and *A. mellifera* (Wu *et al*., 2021). Statistical significance was assessed using Wilcoxon rank-sum tests (two groups) or Kruskal–Wallis tests (≥3 groups), with Benjamini–Hochberg adjustment where applicable. Significance levels: *P* < 0.05 (*); < 0.01 (**); < 0.001 (***); < 0.0001 (****); ns, not significant.

Comparisons between hosts further revealed striking differences in the diversity and abundance of secretion systems. Among young bees, *A. mellifera* exhibited significantly higher secretion system diversity than *A. cerana*, a difference that corresponded to the greater diversity of contributing taxa in *A. mellifera* (Fig. 5B). Consistent with this, certain secretion systems, including elevated T6SS and T9SS, were more abundant in *A. cerana* and were strongly influenced by the *A. cerana*-characteristic taxon *Apibacter* (Fig. 5B, Fig. S14). Interestingly, these host-specific differences diminished with age. In older bees, both the taxonomic composition and secretion system diversity did not differ significantly between *A. mellifera* and *A. cerana* (Fig. 5C).

## Discussion

Here, we show that the secretion systems of the bee gut microbiome follow an evolutionary trajectory distinct from that described in most free-living and pathogenic bacteria. Rather than being shaped by frequent horizontal gene transfer, secretion systems in bee gut microbiome are predominantly vertically inherited and evolve through recurrent gene loss. This pattern reveals secretion systems as long-term ecological traits, retained and maintained due to their advantage in stable host-associated conditions, rather than being employed in transient adaptive responses to fluctuating environments.

Across multiple bacterial lineages of the bee gut microbiome, we observe secretion systems as primarily vertically inherited, indicating long-term co-divergence with their bacteria. Given the prevalence of T1SS, T5SS and T6SS, we infer that interactions within the bee gut niche are heavily driven by interbacterial competition (Kanonenberg et al. 2018; Coulthurst 2019; Meuskens et al. 2019). Theoretical predictions suggests that in such competitive niches, these interaction traits would be subjected to horizontal transfers (Coyte et al. 2015; Niehus et al. 2015). Consistent with these predictions, horizontal transfer of secretion systems is widely reported in other gut-associated and pathogenic bacteria. In the human gut, T6SS loci located on integrative and conjugative elements (ICEs) can spread across co-resident *Bacteroidales* species within and between hosts (Coyne et al. 2014; García-Bayona et al. 2021). Similarly, in the core mouse gut bacteria *Mucispirillum schaedleri*, a T6SS locus appears to have been acquired from distantly related taxa such as *Campylobacter* or *Helicobacter* (Loy et al. 2017). In pathogens such as *Legionella pneumophila* and *Neisseria gonorrhoeae*, the type IV pili and T4SS, both facilitating DNA uptake, are located on large mobile genetic elements and thus allow their own horizontal dissemination (Hamilton et al. 2005; Lautner et al. 2013). More broadly, T3SS and flagella are frequently horizontally acquired across diverse taxa, including environmental *Vibrio cholerae*, plant pathogens such as *Erwinia* and *Dickeya*, and animal pathogens including *Salmonella*, *Shigella*, and *Yersinia* (Naum et al. 2009; Brown and Finlay 2011; Abby and Rocha 2012; Morita et al. 2013).

The contrasting evolutionary trajectory in the bee gut microbiome might be due to the fundamental ecological differences of this environment. The bee gut represents a stable, host-restricted environment in which microbial transmission occurs primarily through social interactions (Powell et al. 2014; Kwong and Moran 2016a). In such a closed system, there is limited exposure to external microbial gene pools and, therefore, the observed vertical inheritance likely reflects strong selection for maintaining interaction traits that are consistently beneficial across host generations. Furthermore, we find that host identity exerts only a weak influence on secretion system repertoires despite the described host specificity among strains (Kwong et al. 2014; Powell et al. 2016; Ellegaard et al. 2020; Zhou et al. 2025). This contrasts with free-living and pathogenic systems, where fluctuating environments and host immune pressures drive an evolutionary arms race and promote horizontal acquisition of invasion and immune-evasion mechanisms (Bliven and Maurelli 2016). Notably, the well-described systems in this context, T2SS and T3SS, also happen to be mostly acquired horizontally (Gophna et al. 2003; Brown and Finlay 2011; Abby and Rocha 2012; Cianciotto and White 2017), and are completely missing in the bee gut microbiome. This absence does not imply an absence of host interaction: several systems retained across bee gut lineages have been described to mediate bacterial-host interactions in other bacteria. Rather, these interactions likely occur within a stable, co-evolved framework, where secretion systems are maintained because they provide long-term persistence advantage. Rare horizontal acquisitions, such as the subtype i1 T6SS in *Frischella*/*Gilliamella* and T4SS in some *Snodgrassella* strains, may represent lineage-specific exceptions, providing adaptive fine-tuning under intense competitive pressure.

We observe gene losses as the dominant evolutionary force shaping secretion system repertoires. Analysis of gain and loss events revealed repeated, independent losses occurring largely within each bee gut microbial lineage, rather than abrupt one-time gain/loss events at the base of these clades. This pattern indicates that, following evolutionary association with bee gut, different strains within the same lineage have repeatedly and irreversibly lost specific secretion systems while remaining viable and persistent members of the community. Thus, these losses do not only imply a uniform drive toward streamlining of secretion system repertoires, likely due to the natural deletion bias in bacteria (Kuo and Ochman 2009), but also point to multiple lineage- and strain-specific strategies for persistence within the gut ecosystem. For example, while systems such as the T6SS and T1SS are widespread within certain lineages, their repeated loss in distinct branches demonstrates that these systems are not universally required for persistence in the bee gut, and that strains can instead adopt alternative ecological strategies to persist (perhaps niche partitioning or reliance on other interaction systems). In this context, the rare lineage-restricted gain events detected, such as the apparent reacquisition of T4SS variants in some *Bartonella* strains, might best be interpreted as a compensatory mechanism for persistence.

Within this evolutionary framework, the secretion systems retained across the bee gut microbiome represent functional solutions to their ecological demands, including colonisation, persistence and competition. Widely distributed across this community are the T1SS and T5SS, the latter being structurally simple yet functionally versatile. In other bacteria, T5aSSs are associated with enzymatic activity and host interaction, T5bSSs with antibacterial competition, and T5cSSs with adhesion, autoaggregation, and biofilm formation (Linke et al. 2006; Aoki et al. 2010; Leo et al. 2010; Serra et al. 2011; Casasanta et al. 2017; Michalska et al. 2017; Trunk et al. 2018; Gucinski et al. 2019; Meuskens et al. 2019; Cuthbert et al. 2022; Qin et al. 2022). Likewise, the T1SS functions as a generalist system mediating biofilm formation, nutrient acquisition, and interbacterial competition through the secretion of adhesins, degradative enzymes, siderophores, and antimicrobials (Thomas et al. 2014; Green and Mecsas 2016; Sassone-Corsi et al. 2016; De León et al. 2017; Kanonenberg et al. 2018; Smith et al. 2018; Mortzfeld et al. 2022; Parker and Davies 2022; Bui et al. 2023; Karbelkar et al. 2024). The high prevalence of these systems and the distinct subtype distribution of the T5SS suggest lineage-specific ecological strategies within the community.

An example of this lineage-specific patterning is observed in *Bartonella* and *Commensalibacter*, which are enriched in T1SS and either the T5aSS or T5bSS while lacking other secretion systems. Although these taxa persist alongside the abundant, multi-system *Frischella*/*Gilliamella* and *Snodgrassella*, they exhibit marked dominance in aged and winter bees, a characteristic shift linked to dietary changes, particularly reduced pollen intake (Kešnerová et al. 2020; C. Li et al. 2022). In this context, reliance on T1SS and T5aSS-associated enzymatic functions may confer a competitive advantage in nutrient-limited gut environments. In *Bartonella*, the presence of two T1SS copies, including one likely acquired early in its evolutionary association with bees, suggests a gain of function that may further enhance persistence and competitive fitness in densely populated niches (Galán 2009; Mendonça et al. 2011; Ghosh and O’Connor 2017). By contrast, in *Commensalibacter*, which encodes only T1SS and T5bSS, these systems might partition roles, with the T1SS mediating secretion of degradative enzymes and adhesins and the T5bSS facilitating competitive interactions. A similar ecological strategy may be observed in *Bombella* (formerly *Saccharibacter* and *Parasaccharibacter*; Fig. S3), which inhabits nutrient-rich niches such as honey and pollen, where T5aSS-mediated enzymatic activity likely enhances access to complex substrates (Zheng et al. 2016; Kwong, Mancenido, et al. 2017; Zheng et al. 2019; Horak et al. 2020). Lastly, the T5cSS, which is mostly restricted to *Snodgrassella*, has been demonstrated to be essential for exporting adhesin to initiate cell-to-cell connections, autoaggregation, and eventually biofilm formation in this lineage (Lariviere et al. 2024).

The T6SS is a contractile nanomachine best known for mediating interbacterial competition, but has also been found to function in nutrient acquisition, spatial and temporal organisation, biofilm formation, and stress resistance (Zhang et al. 2011; Weber et al. 2015; Wong et al. 2016; Lin et al. 2017; Speare et al. 2018; Chen et al. 2019; Custodio et al. 2021; Serapio-Palacios et al. 2022; Xiong et al. 2024; Liang et al. 2025; Luo et al. 2025). In the bee gut microbiome, studies have shown a role for T6SS in mediating interbacterial competition and bacterial-host interactions: in *Snodgrassella*, T6SS subtype i3 mediates interspecific bacterial competition and are associated with Rhs family of toxins (Steele et al. 2017; Steele and Moran 2021), and the subtype i2 is involved in stimulating host immune functions (Motta et al. 2024). Given the close phylogenetic relationship with the T6SS subtype i2 of *Snodgrassella* (Steele et al. 2017), the T6SS subtype i2 in *Gilliamella* may also be involved in bacteria-host interaction, just as reported in *Frischella* (Schmidt et al. 2023). In fact, *Gilliamella* and *Snodgrassella* can exist within a layered biofilm close to each other in the ileum (Martinson et al. 2012), yet reports suggest their loads in the gut are not affected by each other in a T6SS-dependent manner (Motta et al. 2024; Quinn et al. 2024). The T6SS subtype i1 in *Frischella*, which is also horizontally acquired by a subset of *Gilliamella* strains lacking subtype i2, is likely to mediate interbacterial competition; accordingly, its horizontal acquisition may represent an adaptive mechanism promoting competitive fitness and persistence in these strains.

The less prevalent secretion systems of the bee microbiome further support lineage-specific niche specialization. The T4SS is rare and exhibits patchy distribution across members of *Bartonella*, *Bombella*, *Gilliamella*, and *Snodgrassella*. In *Bartonella*, we find an early loss of the system during the diversification of the bee-associated lineage. Perhaps this loss was a prerequisite for the establishment of a non-pathogenic relationship with their bee hosts, since the T4SS in other *Bartonella* spp. has been shown to secrete virulence factors (Schmid et al. 2004; Schröder et al. 2011; Siamer and Dehio 2015). While the apparent re-acquisition in two *Bartonella* strains of a T4SS closely related to the pathogenic *Brucella/Ochrobactrum* seem to contradict this hypothesis, these may represent adaptations restricted to these strains to evade the host immune response during the niche establishment (Dugelay et al. 2025; Wei et al. 2025). On the contrary, the T4SSs either retained or independently acquired in *Bombella*, *Gilliamella*, and *Snodgrassella* correspond to conjugative systems, which primarily mediate DNA transfer and are therefore more likely to function in facilitating horizontal gene exchange within these lineages (Waksman 2025; Zhang et al. 2025). The retention of the T2SS- and T3SS-evolutionary related appendages Tad, T4P, and flagella, further highlights the importance of adhesion, surface sensing, motility, and biofilm formation in lineage-specific niche specialization (Mignolet et al. 2018; Ligthart et al. 2020; Grognot and Taute 2021; Akahoshi and Bevins 2022; Geiger et al. 2024; Whitfield and Brun 2024). Tad and T4P are particularly enriched in *Gilliamella* and *Snodgrassella*, the dominant, co-occurring taxa within the ileum, where their adhesion and biofilm functions to support persistence and facilitate coexistence through spatiotemporal regulation within a confined niche (Y. Li et al. 2022). In lineages lacking Tad and T4P, such as *Bartonella* and *Bombella*, flagella-mediated motility and surface sensing might provide an alternative strategy to locate and colonize available niches.

*Apibacter* represents an extreme case of secretion system streamlining, encoding only T6SS and T9SS. The T9SS, restricted to the *Fibrobacteres–Chlorobi–Bacteroidetes* superphylum, mediates secretion of proteins involved in gliding motility, biofilm formation, macromolecule degradation, metal acquisition, and virulence (Lasica et al. 2017; Veith et al. 2017; Rocha et al. 2024; Niu et al. 2025). Given *Apibacter*’s limited capacity for monosaccharide utilization and unlikely involvement in complex polysaccharide degradation (Kwong and Moran 2016b; Praet et al. 2016; Kwong et al. 2018; W. Zhang et al. 2022), the universal presence of T9SS likely support complementary functions in polysaccharide processing. *Apibacter* primarily colonizes the guts of *Apis cerana*, *Apis dorsata*, and bumble bees, co-occurring with *Snodgrassella* in *A. cerana* but occupying niches where *Snodgrassella* is absent in A. dorsata (Kwong and Moran 2016b; Kwong, Medina, et al. 2017; Kwong et al. 2018; W. Zhang et al. 2022; Suzuki et al. 2025). The presence of T9SS may allow *Apibacter* to utilize host-derived glycans or colonize specific surfaces inaccessible to *Snodgrassella* and other lineages. Although the role of T9SS in interbacterial interactions remains poorly characterized, it has been shown to secrete antibacterial toxins (Jana et al. 2020). In addition, most of the Rhs toxins in *Apibacter* appear to be associated with the T9SS (Steele et al. 2017; Steele and Moran 2021) suggesting a role in bacterial competition. Perhaps, while T9SS mediates nutrients acquisition and bacterial competition, T6SS further enhances fitness by modulating host immunity, potentially providing a selective advantage in certain hosts.

Finally, our metagenomic analysis further underpins how the secretion system repertoires reflect the colonization, persistence and maintenance of the bee gut microbiome as a community. Functionally, elevated Tad, T4P, and T6SS in young bees might indicate that early gut communities are focused on colonization and interbacterial competition, consistent with the initial assembly of the bee gut microbiota, where successful colonizers rely on adhesion, niche acquisition, and competitive persistence (Powell et al. 2016; Jones et al. 2025). In contrast, flagella dominate in older bees, possibly reflecting a shift toward persistence and resource exploitation in gut environment where the hosts’ dietary consumption both changes and declines (Crailsheim et al. 1992). However, we acknowledge that metagenomic inference is inherently limited by its reliance on short-read mapping. Even with our curated bee genera secretion system database, this approach may underestimate diversity, and overlook divergent loci, low-abundance taxa, or strain-level variation absent from reference genomes. In addition, because our analysis is not based on metagenome-assembled genomes, the presence of component genes outside the context of their secretion system architectures does not necessarily indicate a complete system. As such, the observed abundance patterns should be interpreted as approximations of community-level trends rather than direct measures of functional activity.

Taken together, our comparative genomic, evolutionary, and metagenomic analyses converge on a model in which secretion systems in bee gut symbionts are primarily vertically inherited and shaped predominantly by gene losses. At the same time, ecological pressures, including host age, host species, and interbacterial competition, modulate their abundance in vivo. The result is a community that balances evolutionary conservation with ecological plasticity: stable, vertically transmitted systems underpin long-term host association, while dynamic shifts in deployment across host development reinforce functional specialization and gut microbiota stability. Our study establishes a framework for understanding how secretion systems contribute to the stability of the bee gut microbiota and provides a basis for future experimental investigations into how microbial interactions influence community structure and host physiology.

## Materials and methods

### Genomic and metagenomic data

Genomic data were obtained for the core Gram-negative bacterial lineages associated with the bee gut microbiota, including *Apibacter*, *Bartonella*, *Bombella*, *Commensalibacter*, *Gilliamella*, *Frischella*, and *Snodgrassella*. For each lineage, NCBI RefSeq GenBank genome assemblies and corresponding proteomes were downloaded in August 2023. One representative genome from both closely related (same genera) and distantly related (same family) species within each bacterial lineage was included for comparative analyses and for phylogenetic rooting. Tables S1-S5 summarize the bacteria strains, accession numbers, and geographical information associated with the genomic data used in this study. Raw sequencing reads from published bee gut metagenomic datasets (Ellegaard and Engel 2019; Ellegaard et al. 2020; Wu et al. 2021) were retrieved from NCBI using the SRA Toolkit, based on their corresponding BioProject and SRA (SRR) accessions.

### Whole-genome phylogeny

A whole-genome phylogeny for each lineage was constructed based on the concatenated single-copy genes. Single-copy orthologues were inferred from the set of proteomes for each family: *Weeksellaceace* (*Apibacter* and outgroups, 302 orthologues), *Bartonellaceae* (*Bartonella* and outgroups, 247 orthologues), *Acetobacteraceae* (*Bombella* & *Commensalibacter* and outgroups, 24 orthologues), *Orbaceae* (*Frischella* & *Gilliamella* and outgroups, 337 orthologues), and *Neisseriaceae* (*Snodgrassella* and outgroups, 174 orthologues), using the default parameters of OrthoFinder v2.5.4 (Emms and Kelly 2019). The corresponding coding sequences of the orthologues were extracted and aligned based on their codon structure with the *-alignSeq* parameter of MACSE v2.06 (Ranwez et al. 2011; Ranwez et al. 2021). The aligned sequences were concatenated, and maximum likelihood trees were inferred using the substitutional model finder of IQ-Tree v2.2.0.3 (Minh et al. 2020). Based on the model finder module, the trees for *Apibacter*, *Bartonella*, *Bombella* & *Commensalibacter*, and *Gilliamella* & *Frischella* were inferred using general time-reversible (GTR) substitutional models with empirical base frequencies, proportion of invariant sites and a free rate model of rate categories. The tree of the *Snodgrassella* lineage was inferred with the symmetric (SYM) substitutional model. Tree confidences were tested with 1000 bootstrapping replicates. Phylogenetic trees were visualised on Interactive Tree of Life (iTOL) viewer v7 (Letunic and Bork 2024).

### Detection and distribution of bacterial secretion systems in genomes

To detect the secretion systems in the bee gut microbiome, we employed a customized, curated approach rather than relying on the broad, general-purpose genome annotations produced by tools such as Prokka or the NCBI Prokaryotic Genome Annotation Pipeline (PGAP) or relying solely on protein identity scores using tools such as BLAST. By targeting secretion system–specific gene sets and architectures, this approach achieves greater sensitivity and accuracy than both default annotations and previous functional gene surveys of these systems (Barret et al. 2013; Ugarte-Ruiz et al. 2015; Wallner et al. 2021; Liyanapathiranage et al. 2022; Robinson et al. 2022; Robinson et al. 2023).

Components of the type I, II, III, IV, V and IX secretion systems and evolutionarily related appendages tight adherence (Tad), type IV pili (T4P) and flagellum were detected with Macsyfinder TXSScan v1.1.1 (Abby et al. 2016). For each genome, we generated a proteome FASTA file directly from the corresponding GenBank annotation using an in-house script. This approach was preferred over the use of the NCBI-provided proteome files, which include only representative proteins based on protein accession numbers and may omit duplicated proteins. In contrast, our custom-generated proteomes retain proteins according to their locus tags, allowing accurate identification of duplicated or paralogous components. Components of the type VI secretion systems were detected with SecReT6 v3 (J. Li et al. 2015; J. Zhang et al. 2022) which inputs the GenBank file and thus takes into consideration duplicates and genomic locations.

TXSScan detects components of each secretion system using hidden Markov model (HMM) profile-based searches with HMMER (Abby et al. 2016). A system was reported as present based on the maximum allowable distance between genes, the minimum number of mandatory components, and the total number of components detected. Secretion systems are typically composed of multiple proteins, among which a highly conserved core set is essential for assembly and function; these core components are generally encoded together within an operon (Green and Mecsas 2016). This detection strategy leverages the characteristic genomic organisation of secretion systems. Core components, which are highly conserved and essential, form the basis of the computational model and are required for a system to be reported. The less conserved or functionally redundant accessory components are included to improve sensitivity while avoiding false positives. The software evaluates whether detected components are co-localized, reflecting their typical arrangement in operons, and applies thresholds for the minimum number of mandatory components and overall system completeness. Likewise, the genomic organisation is used to further infer systems being encoded in a single locus, where most components occur together, or as multi-loci with components distributed across several genomic regions. We set the maximum number of genes between the components to 5 for T1SS, T2SS, T4P, Tad, and T9SS; 10 for T3SS; 20 for flagellum, and 1 for the subtypes of T5SS. The minimum number out of the total mandatory components was set as 3 out of 3 for T1SS; 3 out of 4 for T2SS; 7 out of 9 for T3SS; 4 out of 6 for T4P; 4 out of 7 for Tad; 9 out of 11 for flagellum; 7 out of 8 for T9SS; and 1 for the subtypes of T5SS. Lastly, the minimum required total number of components was set to 3 out of 3 for T1SS; 6 out of 13 for T2SS; 7 out of 9 for T3SS; 5 out of 18 for T4P; 6 out of 13 for Tad; 10 out of 11 for flagellum; 7 out of 11 for T9SS; and 1 for each subtype of T5SS. These thresholds were chosen to optimize detection sensitivity and minimize false positives for our set of genomes.

SecReT6 v3 was used to detect T6SS components employing a database of experimentally validated protein sequences. Like the TXSScan, this method also considers the minimum number of T6SS components and their genomic organisation. Moreover, phylogenetic analysis of T6SS core components indicates the presence of at least four T6SS subtypes (T6SSi-iv) across different taxa (Boyer et al. 2009). The canonical T6SSi (subdivided into i1, i2, i3, i4a, i4b, and i5) occurs primarily in *Proteobacteria*, T6SSii in *Francisella* pathogenicity islands, T6SSiii in *Bacteriodota*, and T6SSiv in *Amoebophilus* (Boyer et al. 2009; Bröms et al. 2010; Russell et al. 2014; Bayer-Santos et al. 2019). To detect T6SS in the genomes, hits were initially considered valid for components with a protein identity of ≥ 30% to validated components, and a group of co-localized components was regarded as a locus when consisting of ≥ 5 different types of components. The TssB protein, which has been shown to be sufficient to classify the T6SS (J. Li et al. 2015) was extracted and used to build a phylogeny for classification on the SecReT6 web server. Only T6SSi and T6SSiii existed in the datasets. Based on this, a T6SS was categorized as complete/putative when it had ≥ 12 for subtype T6SSiii (disregarding the less well-characterized TssQ and TssR) and ≥ 13 for subtype T6SSi (noting that TssI occurred farther away from the locus in some genomes). The detected proteins were further analysed manually and visually using Geneious Prime v2024.0.7 to identify possible false positives, as previously reported (Steele and Moran 2021), with some modifications. Briefly, a ≥ 50% identity cutoff and a minimum length of 800 amino acids were applied to filter TssH hits, which mostly occurred in loci with two or more detected TssH. An amino acid length of ≥300 was used to filter out TssI hits. T6SSiii lacks TssL; however, at 30% amino acid identity, OmpA-like proteins (Russell et al. 2014; Zoued et al. 2014; Wang et al. 2018) were detected and considered false positives, and were thus removed.

### Phylogenies of secretion systems

Phylogenetic trees were constructed for secretion systems detected in each lineage using the concatenated protein sequences of components that appear as single copy in a locus/region. A system was considered occurring as a locus when the number of unique components as mentioned above co-occurred. In most instances, most, if not all, components co-occurred in a single locus and thus were assigned a locus number based on their genomic coordinates. In some instances, such as the T6SS, set of unique components co-occurred in 2 genomic locations but were considered as a single system as the set of components on either locus constituted the total component count for that system. The phylogenetic trees were constructed based on the components within the locus, either intact or combined.

To this end, T1SS phylogenies for *Bartonella* and *Bombella* & *Commensalibacter* were inferred using concatenated ABC transporter (Abc), membrane fusion protein (Mfp), and outer membrane factor (Omf) proteins, whereas T1SS phylogenies for *Frischella* & *Gilliamella* and *Snodgrassella* were constructed using Mfp and Omf only. Flagellar phylogenies were reconstructed using FlgB, FliE, and the shared T3SS components SctJNRSTUV for *Bartonella*; FlgBC, FliE, and SctJNQRSTUV for *Bombella* & *Commensalibacter*; and FliE together with SctJRSTUV for *Frischella* & *Gilliamella*. Type IV pilus (T4P) phylogenies were inferred using FimT and PilBCDMOQTVW for *Frischella* & *Gilliamella*, and FimT together with PilBCEMNOPQTUVWX for *Snodgrassella*. The tight adherence (Tad) system phylogeny for *Snodgrassella* was reconstructed using concatenated RcpA and TadABCDFV proteins. For *Apibacter*, the T9SS phylogeny was inferred from concatenated GldJKLMN, PorQUV, and SprAET proteins. Finally, T6SS phylogenies were constructed using concatenated TssBCEFGHIKNOP for *Apibacter*, TssABCDEGHJKLM for *Frischella* & *Gilliamella*, and TssABCDEFGHIKLM for *Snodgrassella*.

T6SS phylogenies were constructed in two ways: (i) subtype-specific trees, based on the phylogenetic classification of the conserved TssB protein; and (ii) a combined T6SS (all subtypes) tree, generated by concatenating all single-copy T6SS loci present in each genome without separating subtypes into independent trees. In this combined tree, loci belonging to different T6SS subtypes remain distinct concatenated components in the alignment but are included together in a single phylogeny. The locus tags of each component were used to extract their corresponding protein sequences. The sequences were aligned using MUSCLE v5.1.linux64 (Edgar 2022), concatenated, and phylogenetic trees were constructed using IQ-Tree v2.2.0.3 (Minh et al. 2020). The trees for each lineage were constructed with the LG+F+I+G4 model and tested with 1000 bootstrap replicates. For each system, the individual component gene trees were also inferred under the same model with ultrafast booststrap distributions with 1000 bootstrap replicates. Phylogenetic trees were visualized on iTOL v7 (Letunic and Bork 2024).

### Test for phylogenetic congruence and reconciliation

The co-evolutionary relationships between secretions systems and their bacteria were tested with three separate distance-based cophylogenetic methods, ParaFit (Legendre et al. 2002), Hommola (Hommola et al. 2009), and PACo (Hutchinson et al. 2017), and one event-based method, eMPRess (Santichaivekin et al. 2021). These methods were originally described to test for co-speciation between parasites and their hosts by fitting their respective phylogenetic distances.

Briefly, these pattern-based methods use the secretion system (treated as the parasite) and lineage (treated as the host) phylogenetic distances to test whether an interaction between the two phylogenies is due to chance alone. Using the R script from (Nishida and Ochman 2021), the phylogenetic distance matrices were computed from each phylogenetic tree, transformed into principal coordinates and superimposed to assess the degree of evolutionary congruence while accounting for lineage-secretion system associations. Each lineage–secretion system pair was analysed through 10 simulations, with 999 permutations performed for each simulation. The tests were performed as Type I error tests with the null hypothesis that the lineage and their secretion systems evolved independently, i.e., a *p*-value ≤ 0.05 signifies congruence between both phylogenies.

The event-based method reconciles the two phylogenies by mapping the system phylogeny onto the bacterial lineage phylogeny. Reconciliations were performed with the loss cost fixed at 1, while duplication and transfer costs were independently varied from 1 to 3, generating nine cost combinations in total. For each cost scheme, a reconciliation using maximum parsimony was computed and statistical significance was assessed with 100 samplings using the built-in randomization (*p*-value) module. The resulting *p*-values from the nine analyses were averaged to obtain a single summary estimate of cophylogenetic significance.

### Metagenome assembly and annotation

Publicly available raw metagenomic data and metadata (Ellegaard and Engel 2019; Ellegaard et al. 2020; Wu et al. 2021) were prefetched from the NCBI SRR repository. These are metagenomes obtained from either *Apis mellifera* or *Apis cerana*. For each metagenomic sample, paired-end reads were extracted, and the quality of the raw reads was checked with fastQC v0.12.0. Since each metagenome was sequenced with different technologies, Trim Galore v0.6.10 was used to trim the reads using the auto default settings, which include an initial quality check before trimming detected the adaptor. QC logs were manually inspected to ensure the trimmed adaptors corresponded with the reported adaptors in their source studies. Contaminating host-derived reads were filtered out by first mapping the raw reads against a database of the source host genomes, *A. mellifera* (PRJNA471592) or *A. cerana* (PRJNA324433), using Bowtie2 v2.5.3 (Langmead and Salzberg 2012). Samtools v1.19.2 (Danecek et al. 2021) was used to generate bam-files of the filtered reads (*view -b -f 12 -F 256*), sort, and extract the paired-end reads. Each pair of unfiltered and filtered reads was profiled for their represented communities using Kraken2 v2.1.3 (Wood et al. 2019), visualized as a community profile and manually inspected to validate the removal of host-derived reads. The filtered paired-end reads were assembled using the default setting of MEGAHIT assembler v1.2.9 (D. Li et al. 2015). The resulting contigs were filtered to remove contigs shorter than 1000 bp. Protein coding sequences of the contigs were predicted using Prokka v1.14.6 (Seemann 2014).

### Detection and distribution of secretion systems in metagenome data

Secretion system components were identified in metagenomic assemblies based on protein sequence similarity. A custom BLAST database was constructed using all predicted protein sequences of secretion system components from our individual genome analysis, with each protein linked to its corresponding genome metadata. Predicted protein sequences from the metagenomes were queried against this database using BLASTp v 2.12.0+ (e-value < 1e-20, minimum identity > 40%), retaining only the top hit for each query. To distinguish bee-specific components, we examined the distribution of sequence identities for each component and established thresholds based on modal distributions. A minimum threshold of ≥70% identity was required for considering a component as bee specific. Coding sequences corresponding to each component were further taxonomically classified using Kraken2 v2.1.3 (Wood et al. 2019) and validated manually through BLASTn searches against the NCBI non-redundant nucleotide database. Manual inspection was performed to confirm that the classified genus of contigs matched the genus of the corresponding reference protein, supporting their assignment as bee-associated bacterial components. Potential false positives were noted for some components, particularly ATPases of the type III secretion system (T3SS) and flagellar system, which occasionally occurred as singletons in taxa where no complete system was detected in genome analyses. To avoid this, we required at least two unique components for a secretion system to be considered present in each metagenome, removing hits based solely on single components. Exceptions to this were the T5SS, where the subtypes are single component systems.

The relative abundance of each secretion system was estimated from the summed abundances of its constituent components. All predicted coding sequences were mapped against quality-filtered metagenomic reads using BWA-MEM2 v2.2.1 (Vasimuddin et al. 2019). Coverage was calculated with BEDTools v2.31.1 (Quinlan and Hall 2010), and read counts were normalized by gene length. To account for differences in sequencing depth across samples, abundances were further normalized as copies per million reads (CPM). From the overall abundance matrix, we extracted secretion system components to calculate (i) the total abundance of each secretion system per metagenome, and (ii) diversity indices of secretion systems within and across samples.

## Author contributions

**Samuel A. Acheampong**: Conceptualization; Data curation; Software; Formal analysis; Validation; Investigation; Visualization; Methodology; Writing—original draft; Writing—review and editing. **Waldan K. Kwong**: Conceptualization; Resources; Supervision; Funding acquisition; Validation; Writing—original draft; Project administration; Writing—review and editing.

## Acknowledgements

The authors thank members of the Microbial Genomics and Symbiosis lab for constructive discussions, and Sara Louro for assistance in curating metagenome data. This work was supported by Fundação para a Ciência e a Tecnologia (FCT) doctoral fellowship UI/BD/154902/2023 to SAA, and European Molecular Biology Organization Installation Grant 5045-2022, European Research Council Starting Grant 101042912, and FCT individual grant CEECIND/01358/2021 to WKK.

## References

Abby SS, Cury J, Guglielmini J, Néron B, Touchon M, Rocha EPC. 2016. Identification of protein secretion systems in bacterial genomes. Sci. Rep. 6:23080.

Abby SS, Rocha EPC. 2012. The non-flagellar type III secretion system evolved from the bacterial flagellum and diversified into host-cell adapted systems. PLOS Genet. 8:e1002983.

Akahoshi DT, Bevins CL. 2022. Flagella at the host-microbe Interface: key functions intersect with redundant responses. Front. Immunol. 13:828758.

Aoki SK, Diner EJ, de Roodenbeke C t’Kint, Burgess BR, Poole SJ, Braaten BA, Jones AM, Webb JS, Hayes CS, Cotter PA, et al. 2010. A widespread family of polymorphic contact-dependent toxin delivery systems in bacteria. Nature 468:439–442.

Baishya J, Wakeman CA. 2019. Selective pressures during chronic infection drive microbial competition and cooperation. Npj Biofilms Microbiomes 5:16.

Barret M, Egan F, O’Gara F. 2013. Distribution and diversity of bacterial secretion systems across metagenomic datasets. Environ. Microbiol. Rep. 5:117–126.

Bayer-Santos E, Ceseti L de M, Farah CS, Alvarez-Martinez CE. 2019. Distribution, function and regulation of type 6 secretion systems of *Xanthomonadales*. Front. Microbiol. 10:1635.

Bliven KA, Maurelli AT. 2016. Evolution of bacterial pathogens within the human host. Microbiol. Spectr. 4:10.1128/microbiolspec.vmbf-0017-2015.

Bontemps Z, Paranjape K, Guy L. 2024. Host–bacteria interactions: ecological and evolutionary insights from ancient, professional endosymbionts. FEMS Microbiol. Rev. 48:fuae021.

Boyer F, Fichant G, Berthod J, Vandenbrouck Y, Attree I. 2009. Dissecting the bacterial type VI secretion system by a genome wide in silico analysis: what can be learned from available microbial genomic resources? BMC Genomics 10:104.

Bröms J, Sjöstedt A, Lavander M. 2010. The role of the *Francisella Tularensis* pathogenicity island in type VI secretion, intracellular survival, and modulation of host cell signaling. Front. Microbiol. 1.

Brown NF, Finlay B. 2011. Potential origins and horizontal transfer of type III secretion systems and effectors. Mob. Genet. Elem. 1:118–121.

Bui D-C, Luo T, McBride JW. 2023. Type 1 secretion system and effectors in *Rickettsiales*. Front. Cell. Infect. Microbiol. 13:1175688.

Büttner D, He SY. 2009. Type III protein secretion in plant pathogenic bacteria. Plant Physiol. 150:1656–1664.

Casasanta MA, Yoo CC, Smith HB, Duncan AJ, Cochrane K, Varano AC, Allen-Vercoe E, Slade DJ. 2017. A chemical and biological toolbox for Type Vd secretion: characterization of the phospholipase A1 autotransporter FplA from *Fusobacterium nucleatum*. J. Biol. Chem. 292:20240–20254.

Chen C, Yang X, Shen X. 2019. Confirmed and potential roles of bacterial T6SSs in the intestinal ecosystem. Front. Microbiol. 10.

Christian N, Whitaker BK, Clay K. 2015. Microbiomes: unifying animal and plant systems through the lens of community ecology theory. Front. Microbiol. 6:869.

Cianciotto NP, White RC. 2017. Expanding role of type II secretion in bacterial pathogenesis and beyond. Infect. Immun. 85:10.1128/iai.00014-17.

Coulthurst S. 2019. The Type VI secretion system: a versatile bacterial weapon. Microbiology 165:503–515.

Coyne MJ, Zitomersky NL, McGuire AM, Earl AM, Comstock LE. 2014. Evidence of extensive DNA transfer between *Bacteroidales* species within the human gut. mBio 5:10.1128/mbio.01305-14.

Coyte KZ, Schluter J, Foster KR. 2015. The ecology of the microbiome: networks, competition, and stability. Science 350:663–666.

Crailsheim K, Schneider LHW, Hrassnigg N, Bühlmann G, Brosch U, Gmeinbauer R, Schöffmann B. 1992. Pollen consumption and utilization in worker honeybees (*Apis mellifera carnica*): dependence on individual age and function. J. Insect Physiol. 38:409–419.

Custodio R, Ford RM, Ellison CJ, Liu G, Mickute G, Tang CM, Exley RM. 2021. Type VI secretion system killing by commensal Neisseria is influenced by expression of type four pili.Filloux A, Storz G, Filloux A, editors. eLife 10:e63755.

Cuthbert BJ, Hayes CS, Goulding CW. 2022. Functional and structural diversity of bacterial contact-dependent growth inhibition effectors. Front. Mol. Biosci. 9:866854.

Danecek P, Bonfield JK, Liddle J, Marshall J, Ohan V, Pollard MO, Whitwham A, Keane T, McCarthy SA, Davies RM, et al. 2021. Twelve years of SAMtools and BCFtools. GigaScience 10:giab008.

De León KB, Zane GM, Trotter VV, Krantz GP, Arkin AP, Butland GP, Walian PJ, Fields MW, Wall JD. 2017. Unintended laboratory-driven evolution reveals genetic requirements for biofilm formation by *Desulfovibrio vulgaris Hildenborough*. mBio 8:10.1128/mbio.01696-17.

Deakin WJ, Broughton WJ. 2009. Symbiotic use of pathogenic strategies: rhizobial protein secretion systems. Nat. Rev. Microbiol. 7:312–320.

Denise R, Abby SS, Rocha EPC. 2020. The evolution of protein secretion systems by co-option and tinkering of cellular machineries. Trends Microbiol. 28:372–386.

Dugelay C, Celli J, Terradot L. 2025. The *Brucella* type IV secretion system and effector proteins. microLife 6:uqaf036.

Edgar RC. 2022. Muscle5: High-accuracy alignment ensembles enable unbiased assessments of sequence homology and phylogeny. Nat. Commun. 13:6968.

Ellegaard KM, Engel P. 2019. Genomic diversity landscape of the honey bee gut microbiota. Nat. Commun. 10:446.

Ellegaard KM, Suenami S, Miyazaki R, Engel P. 2020. Vast differences in strain-level diversity in the gut microbiota of two closely related honey bee species. Curr. Biol. 30:2520–2531.e7.

Emms DM, Kelly S. 2019. OrthoFinder: phylogenetic orthology inference for comparative genomics. Genome Biol. 20:238.

Galán JE. 2009. Common themes in the design and function of bacterial effectors. Cell Host Microbe 5:571–579.

Ganesan R, Wierz JC, Kaltenpoth M, Flórez LV. 2022. How it all begins: bacterial factors mediating the colonization of invertebrate hosts by beneficial symbionts. Microbiol. Mol. Biol. Rev. 86:e00126–21.

García-Bayona L, Coyne MJ, Comstock LE. 2021. Mobile type VI secretion system loci of the gut Bacteroidales display extensive intra-ecosystem transfer, multi-species spread and geographical clustering. PLOS Genet. 17:e1009541.

Geiger CJ, Wong GCL, O’Toole GA. 2024. A bacterial sense of touch: T4P retraction motor as a means of surface sensing by *Pseudomonas aeruginosa* PA14. J. Bacteriol. 206:e00442–23.

Ghosh S, O’Connor TJ. 2017. Beyond paralogs: the multiple layers of redundancy in bacterial pathogenesis. Front. Cell. Infect. Microbiol. 7:467.

Ghoul M, Mitri S. 2016. The ecology and evolution of microbial competition. Trends Microbiol. 24:833–845.

Gophna U, Ron EZ, Graur D. 2003. Bacterial type III secretion systems are ancient and evolved by multiple horizontal-transfer events. Gene 312:151–163.

Gorter FA, Manhart M, Ackermann M. 2020. Understanding the evolution of interspecies interactions in microbial communities. Philos. Trans. R. Soc. B Biol. Sci. 375:20190256.

Green ER, Mecsas J. 2016. Bacterial secretion systems: an overview. Microbiol. Spectr. 4:10.1128/microbiolspec.vmbf-0012-2015.

Grognot M, Taute KM. 2021. More than propellers: how flagella shape bacterial motility behaviors. Curr. Opin. Microbiol. 61:73–81.

Gucinski GC, Michalska K, Garza-Sánchez F, Eschenfeldt WH, Stols L, Nguyen JY, Goulding CW, Joachimiak A, Hayes CS. 2019. Convergent evolution of the Barnase/EndoU/Colicin/RelE (BECR) fold in antibacterial tRNase toxins. Structure 27:1660–1674.e5.

Hamilton HL, Domínguez NM, Schwartz KJ, Hackett KT, Dillard JP. 2005. *Neisseria gonorrhoeae* secretes chromosomal DNA via a novel type IV secretion system. Mol. Microbiol. 55:1704–1721.

Harms A, Liesch M, Körner J, Québatte M, Engel P, Dehio C. 2017. A bacterial toxin-antitoxin module is the origin of inter-bacterial and inter-kingdom effectors of *Bartonella*. PLOS Genet. 13:e1007077.

Hommola K, Smith JE, Qiu Y, Gilks WR. 2009. A permutation test of host–parasite cospeciation. Mol. Biol. Evol. 26:1457–1468.

Horak RD, Leonard SP, Moran NA. 2020. Symbionts shape host innate immunity in honeybees. Proc. R. Soc. B Biol. Sci. 287:20201184.

Hou K, Wu Z-X, Chen X-Y, Wang J-Q, Zhang D, Xiao C, Zhu D, Koya JB, Wei L, Li J, et al. 2022. Microbiota in health and diseases. Signal Transduct. Target. Ther. 7:135.

Hutchinson MC, Cagua EF, Balbuena JA, Stouffer DB, Poisot T. 2017. paco: implementing Procrustean approach to cophylogeny in R. Methods Ecol. Evol. 8:932–940.

Jamali H, Akrami F, Layeghkhavidaki H, Bouakkaz S. 2025. Bacterial protein secretion systems: mechanisms, functions, and roles in virulence. Microb. Pathog. 206:107790.

Jana B, Keppel K, Fridman CM, Bosis E, Salomon D. 2022. Multiple T6SSs, mobile auxiliary modules, and effectors revealed in a systematic snalysis of the Vibrio parahaemolyticus pan-genome. mSystems 7:e00723–22.

Jana B, Salomon D, Bosis E. 2020. A novel class of polymorphic toxins in *Bacteroidetes*. Life Sci. Alliance 3:e201900631.

Jones KR, Song Y, Rinaldi SS, Moran NA. 2025. Effects of priority on strain-level composition of the honey bee gut community. Appl. Environ. Microbiol. 91:e00828–25.

Kanonenberg K, Spitz O, Erenburg IN, Beer T, Schmitt L. 2018. Type I secretion system—it takes three and a substrate. FEMS Microbiol. Lett. 365:fny094.

Karbelkar AA, Font M, Smith TJ, Sondermann H, O’Toole GA. 2024. Reconstitution of a biofilm adhesin system from a sulfate-reducing bacterium in *Pseudomonas fluorescens*. Proc. Natl. Acad. Sci. 121:e2320410121.

Kešnerová L, Emery O, Troilo M, Liberti J, Erkosar B, Engel P. 2020. Gut microbiota structure differs between honeybees in winter and summer. ISME J. 14:801–814.

Kešnerová L, Mars RAT, Ellegaard KM, Troilo M, Sauer U, Engel P. 2017. Disentangling metabolic functions of bacteria in the honey bee gut. PLOS Biol. 15:e2003467.

Kuo C-H, Ochman H. 2009. Deletional Bias across the Three Domains of Life. Genome Biol. Evol. 1:145–152.

Kwong WK, Engel P, Koch H, Moran NA. 2014. Genomics and host specialization of honey bee and bumble bee gut symbionts. Proc. Natl. Acad. Sci. 111:11509–11514.

Kwong WK, Mancenido AL, Moran NA. 2017. Immune system stimulation by the native gut microbiota of honey bees. R. Soc. Open Sci. 4:170003.

Kwong WK, Medina LA, Koch H, Sing K-W, Soh EJY, Ascher JS, Jaffé R, Moran NA. 2017. Dynamic microbiome evolution in social bees. Sci. Adv. 3:e1600513.

Kwong WK, Moran NA. 2016a. Gut microbial communities of social bees. Nat. Rev. Microbiol. 14:374–384.

Kwong WK, Moran NA. 2016b. Apibacter adventoris gen. nov., sp. nov., a member of the phylum Bacteroidetes isolated from honey bees. Int. J. Syst. Evol. Microbiol. 66:1323–1329.

Kwong WK, Steele MI, Moran NA. 2018. Genome Sequences of Apibacter spp., Gut Symbionts of Asian Honey Bees. Genome Biol. Evol. 10:1174–1179.

Langmead B, Salzberg SL. 2012. Fast gapped-read alignment with Bowtie 2. Nat. Methods 9:357–359.

Lariviere PJ, Ashraf AHMZ, Gifford I, Tanguma SL, Barrick JE, Moran NA. 2024. Virulence-linked adhesin drives mutualist colonization of the bee gut via biofilm formation. :2024.10.14.618124. Available from: https://www.biorxiv.org/content/10.1101/2024.10.14.618124v1

Lasica AM, Ksiazek M, Madej M, Potempa J. 2017. The type IX secretion system (T9SS): highlights and recent insights into its structure and function. Front. Cell. Infect. Microbiol. 7:215.

Lautner M, Schunder E, Herrmann V, Heuner K. 2013. Regulation, integrase-dependent excision, and horizontal transfer of genomic islands in *Legionella pneumophila*. J. Bacteriol. 195:1583–1597.

Lee FJ, Miller KI, McKinlay JB, Newton ILG. 2018. Differential carbohydrate utilization and organic acid production by honey bee symbionts. FEMS Microbiol. Ecol. 94:fiy113.

Legendre P, Desdevises Y, Bazin E. 2002. A statistical test for host–parasite coevolution. Syst. Biol. 51:217–234.

Leo JC, Elovaara H, Bihan D, Pugh N, Kilpinen SK, Raynal N, Skurnik M, Farndale RW, Goldman A. 2010. First analysis of a bacterial collagen-binding protein with collagen toolkits: promiscuous binding of YadA to collagens may explain how YadA interferes with host processes. Infect. Immun. 78:3226–3236.

Letunic I, Bork P. 2024. Interactive Tree of Life (iTOL) v6: recent updates to the phylogenetic tree display and annotation tool. Nucleic Acids Res. 52:W78–W82.

Li C, Tang M, Li X, Zhou X. 2022. Community dynamics in structure and function of honey bee gut bacteria in response to winter dietary shift. mBio 13:e01131–22.

Li D, Liu C-M, Luo R, Sadakane K, Lam T-W. 2015. MEGAHIT: an ultra-fast single-node solution for large and complex metagenomics assembly via succinct de Bruijn graph. Bioinforma. Oxf. Engl. 31:1674–1676.

Li DY, Liu YL, Liao XJ, He TT, Sun SS, Nie P, Xie HX. 2021. Identification and characterization of EvpQ, a novel T6SS effector encoded on a mobile genetic element in *Edwardsiella piscicida*. Front. Microbiol. 12.

Li J, Yao Y, Xu HH, Hao L, Deng Z, Rajakumar K, Ou H-Y. 2015. SecReT6: a web-based resource for type VI secretion systems found in bacteria. Environ. Microbiol. 17:2196–2202.

Li Y, Leonard SP, Powell JE, Moran NA. 2022. Species divergence in gut-restricted bacteria of social bees. Proc. Natl. Acad. Sci. 119:e2115013119.

Li YG, Hu B, Christie PJ. 2019. Biological and structural diversity of type IV secretion systems. Microbiol. Spectr. 7:10.1128/microbiolspec.psib-0012-2018.

Liang C, Lin L, Wang X, Zhu W. 2025. The secretion of *Pseudomonas* unconventional peroxidase facilitates extracellular carbon acquisition from heterogeneous lignin. *Commun*. Biol. 8:1318.

Ligthart K, Belzer C, de Vos WM, Tytgat HLP. 2020. Bridging bacteria and the gut: functional aspects of type IV pili. Trends Microbiol. 28:340–348.

Lin J, Zhang W, Cheng J, Yang X, Zhu K, Wang Y, Wei G, Qian P-Y, Luo Z-Q, Shen X. 2017. A *Pseudomonas* T6SS effector recruits PQS-containing outer membrane vesicles for iron acquisition. Nat. Commun. 8:14888.

Linke D, Riess T, Autenrieth IB, Lupas A, Kempf VAJ. 2006. Trimeric autotransporter adhesins: variable structure, common function. Trends Microbiol. 14:264–270.

Liyanapathiranage P, Wagner N, Avram O, Pupko T, Potnis N. 2022. Phylogenetic distribution and evolution of type VI secretion system in the genus *Xanthomonas*. Front. Microbiol. 13:840308.

Loy A, Pfann C, Steinberger M, Hanson B, Herp S, Brugiroux S, Gomes Neto JC, Boekschoten MV, Schwab C, Urich T, et al. 2017. Lifestyle and horizontal gene transfer-mediated evolution of *Mucispirillum schaedleri*, a core member of the murine gut microbiota. mSystems 2:10.1128/msystems.00171-16.

Luo C, Gu H, Pan D, Zhao Y, Zheng A, Zhu H, Zhang C, Li C, Zhang J, Chen C, et al. 2025. *Pseudomonas aeruginosa* T6SS secretes an oxygen-binding hemerythrin to facilitate competitive growth under microaerobic conditions. Microbiol. Res. 293:128052.

Maphosa S, Moleleki LN, Motaung TE. 2023. Bacterial secretion system functions: evidence of interactions and downstream implications. Microbiology 169:001326.

Martinson VG, Moy J, Moran NA. 2012. Establishment of characteristic gut bacteria during development of the honeybee worker. Appl. Environ. Microbiol. 78:2830–2840.

McFall-Ngai M, Hadfield MG, Bosch TCG, Carey HV, Domazet-Lošo T, Douglas AE, Dubilier N, Eberl G, Fukami T, Gilbert SF, et al. 2013. Animals in a bacterial world, a new imperative for the life sciences. Proc. Natl. Acad. Sci. 110:3229–3236.

Mendonça AG, Alves RJ, Pereira-Leal JB. 2011. Loss of genetic redundancy in reductive genome evolution. PLoS Comput. Biol. 7:e1001082.

Meuskens I, Saragliadis A, Leo JC, Linke D. 2019. Type V secretion systems: an overview of passenger domain functions. Front. Microbiol. 10:1163.

Michalska K, Gucinski GC, Garza-Sánchez F, Johnson PM, Stols LM, Eschenfeldt WH, Babnigg G, Low DA, Goulding CW, Joachimiak A, et al. 2017. Structure of a novel antibacterial toxin that exploits elongation factor Tu to cleave specific transfer RNAs. Nucleic Acids Res. 45:10306–10320.

Mignolet J, Panis G, Viollier PH. 2018. More than a Tad: spatiotemporal control of *Caulobacter* pili. Curr. Opin. Microbiol. 42:79–86.

Minh BQ, Schmidt HA, Chernomor O, Schrempf D, Woodhams MD, von Haeseler A, Lanfear R. 2020. IQ-TREE 2: new models and efficient methods for phylogenetic inference in the genomic era. Mol. Biol. Evol. 37:1530–1534.

Moran NA, McCutcheon JP, Nakabachi A. 2008. Genomics and evolution of heritable bacterial symbionts. Annu. Rev. Genet. 42:165–190.

Morita M, Yamamoto S, Hiyoshi H, Kodama T, Okura M, Arakawa E, Alam M, Ohnishi M, Izumiya H, Watanabe H. 2013. Horizontal gene transfer of a genetic island encoding a type III secretion system distributed in *Vibrio cholerae*. Microbiol. Immunol. 57:334–339.

Mortzfeld BM, Palmer JD, Bhattarai SK, Dupre HL, Mercado-Lubio R, Silby MW, Bang C, McCormick BA, Bucci V. 2022. Microcin MccI47 selectively inhibits enteric bacteria and reduces carbapenem-resistant *Klebsiella pneumoniae* colonization in vivo when administered via an engineered live biotherapeutic. Gut Microbes 14:2127633.

Motta EVS, Lariviere PJ, Jones KR, Song Y, Moran NA. 2024. Type VI secretion systems promote intraspecific competition and host interactions in a bee gut symbiont. Proc. Natl. Acad. Sci. 121:e2414882121.

Motta EVS, Moran NA. 2024. The honeybee microbiota and its impact on health and disease. Nat. Rev. Microbiol. 22:122–137.

Naum M, Brown EW, Mason-Gamer RJ. 2009. Phylogenetic evidence for extensive horizontal gene transfer of type III secretion system genes among enterobacterial plant pathogens. Microbiology 155:3187–3199.

Nazir R, Mazurier S, Yang P, Lemanceau P, van Elsas JD. 2017. The ecological role of type three secretion systems in the interaction of bacteria with fungi in soil and related habitats is diverse and context-dependent. Front. Microbiol. 8.

Nicholson KR, Champion PA. 2022. Bacterial secretion systems: Networks of pathogenic regulation and adaptation in mycobacteria and beyond. PLoS Pathog. 18:e1010610.

Niehus R, Mitri S, Fletcher AG, Foster KR. 2015. Migration and horizontal gene transfer divide microbial genomes into multiple niches. Nat. Commun. 6:8924.

Nishida AH, Ochman H. 2021. Captivity and the co-diversification of great ape microbiomes. Nat. Commun. 12:5632.

Niu P, Guo R, Ding C, Yu S. 2025. Advances in the type IX secretion system: an exploration focusing on *Riemerella anatipestifer*. Curr. Res. Microb. Sci. 9:100404.

Okazaki S, Kaneko T, Sato S, Saeki K. 2013. Hijacking of leguminous nodulation signaling by the rhizobial type III secretion system. Proc. Natl. Acad. Sci. 110:17131–17136.

Parker JK, Davies BW. 2022. Microcins reveal natural mechanisms of bacterial manipulation to inform therapeutic development. Microbiology 168:001175.

Peabody CR, Chung YJ, Yen M-R, Vidal-Ingigliardi D, Pugsley AP, Saier MH. 2003. Type II protein secretion and its relationship to bacterial type IV pili and archaeal flagella. Microbiology 149:3051–3072.

Perreau J, Moran NA. 2022. Genetic innovations in animal–microbe symbioses. Nat. Rev. Genet. 23:23–39.

Powell E, Ratnayeke N, Moran NA. 2016. Strain diversity and host specificity in a specialized gut symbiont of honeybees and bumblebees. Mol. Ecol. 25:4461–4471.

Powell JE, Martinson VG, Urban-Mead K, Moran NA. 2014. Routes of acquisition of the gut microbiota of the honey bee *Apis mellifera*. Appl. Environ. Microbiol. 80:7378–7387.

Praet J, Aerts M, Brandt ED, Meeus I, Smagghe G, Vandamme P. 2016. Apibacter mensalis sp. nov.: a rare member of the bumblebee gut microbiota. Int. J. Syst. Evol. Microbiol. 66:1645–1651.

Qin J, Hong Y, Morona R, Totsika M. 2022. Cysteine-dependent conformational heterogeneity of *Shigella flexneri* autotransporter IcsA and implications of its function. Microbiol. Spectr. 10:e03410–22.

Quinlan AR, Hall IM. 2010. BEDTools: a flexible suite of utilities for comparing genomic features. Bioinformatics 26:841–842.

Quinn A, El Chazli Y, Escrig S, Daraspe J, Neuschwander N, McNally A, Genoud C, Meibom A, Engel P. 2024. Host-derived organic acids enable gut colonization of the honey bee symbiont Snodgrassella alvi. Nat. Microbiol. 9:477–489.

Ranwez V, Chantret N, Delsuc F. 2021. Aligning protein-coding nucleotide sequences with MACSE. In: Katoh K, editor. Multiple Sequence Alignment: Methods and Protocols. New York, NY: Springer US. p. 51–70. Available from: 10.1007/978-1-0716-1036-7_4

Ranwez V, Harispe S, Delsuc F, Douzery EJP. 2011. MACSE: multiple alignment of coding sequences accounting for frameshifts and stop codons. PLOS ONE 6:e22594.

Raymann K, Moran NA. 2018. The role of the gut microbiome in health and disease of adult honey bee workers. Curr. Opin. Insect Sci. 26:97–104.

Robinson L, Liaw J, Omole Z, Corcionivoschi N, Hachani A, Gundogdu O. 2022. In silico investigation of the genus *Campylobacter* type VI secretion system reveals genetic diversity in organization and putative effectors. *Microb*. Genomics 8:000898.

Robinson LA, Collins ACZ, Murphy RA, Davies JC, Allsopp LP. 2023. Diversity and prevalence of type VI secretion system effectors in clinical *Pseudomonas aeruginosa* isolates. Front. Microbiol. 13.

Rocha ST, Shah DD, Shrivastava A. 2024. Ecological, beneficial, and pathogenic functions of the type 9 secretion system. Microb. Biotechnol. 17:e14516.

Russell AB, Wexler AG, Harding BN, Whitney JC, Bohn AJ, Goo YA, Tran BQ, Barry NA, Zheng H, Peterson SB, et al. 2014. A type VI secretion-related pathway in *Bacteroidetes* mediates interbacterial antagonism. Cell Host Microbe 16:227–236.

Santichaivekin S, Yang Q, Liu J, Mawhorter R, Jiang J, Wesley T, Wu Y-C, Libeskind-Hadas R. 2021. eMPRess: a systematic cophylogeny reconciliation tool. Bioinformatics 37:2481–2482.

Sassone-Corsi M, Nuccio S-P, Liu H, Hernandez D, Vu CT, Takahashi AA, Edwards RA, Raffatellu M. 2016. Microcins mediate competition among *Enterobacteriaceae* in the inflamed gut. Nature 540:280–283.

Schepers MJ, Yelland JN, Moran NA, Taylor DW. 2021. Isolation of the *Buchnera aphidicola* flagellum basal body complexes from the *Buchnera* membrane. PLOS ONE 16:e0245710.

Schmid MC, Schulein R, Dehio M, Denecker G, Carena I, Dehio C. 2004. The VirB type IV secretion system of *Bartonella henselae* mediates invasion, proinflammatory activation and antiapoptotic protection of endothelial cells. Mol. Microbiol. 52:81–92.

Schmidt K, Santos-Matos G, Leopold-Messer S, El Chazli Y, Emery O, Steiner T, Piel J, Engel P. 2023. Integration host factor regulates colonization factors in the bee gut symbiont Frischella perrara.Xavier KB, Garrett WS, Xavier KB, Leulier F, Florez Platino L, editors. eLife 12:e76182.

Schröder G, Schuelein R, Quebatte M, Dehio C. 2011. Conjugative DNA transfer into human cells by the VirB/VirD4 type IV secretion system of the bacterial pathogen *Bartonella henselae*. Proc. Natl. Acad. Sci. 108:14643–14648.

Seemann T. 2014. Prokka: rapid prokaryotic genome annotation. Bioinformatics 30:2068–2069.

Serapio-Palacios A, Woodward SE, Vogt SL, Deng W, Creus-Cuadros A, Huus KE, Cirstea M, Gerrie M, Barcik W, Yu H, et al. 2022. Type VI secretion systems of pathogenic and commensal bacteria mediate niche occupancy in the gut. Cell Rep. [Internet] 39. Available from: https://www.cell.com/cell-reports/abstract/S2211-1247(22)00492-2

Serra DO, Conover MS, Arnal L, Sloan GP, Rodriguez ME, Yantorno OM, Deora R. 2011. FHA-mediated cell-substrate and cell-cell adhesions are critical for *Bordetella pertussis* biofilm formation on abiotic surfaces and in the mouse nose and the trachea. PLOS ONE 6:e28811.

Siamer S, Dehio C. 2015. New insights into the role of *Bartonella* effector proteins in pathogenesis. Curr. Opin. Microbiol. 23:80–85.

Smith TJ, Font ME, Kelly CM, Sondermann H, O’Toole GA. 2018. An N-terminal retention module anchors the giant adhesin LapA of *Pseudomonas fluorescens* at the cell surface: a novel subfamily of type I secretion systems. J. Bacteriol. 200:e00734–17.

Speare L, Cecere AG, Guckes KR, Smith S, Wollenberg MS, Mandel MJ, Miyashiro T, Septer AN. 2018. Bacterial symbionts use a type VI secretion system to eliminate competitors in their natural host. Proc. Natl. Acad. Sci. 115:E8528–E8537.

Spitz O, Erenburg IN, Beer T, Kanonenberg K, Holland IB, Schmitt L. 2019. Type I secretion systems—one mechanism for all? Microbiol. Spectr. 7:10.1128/microbiolspec.psib-0003-2018.

Steele MI, Kwong WK, Whiteley M, Moran NA. 2017. Diversification of Type VI Secretion System Toxins Reveals Ancient Antagonism among Bee Gut Microbes. mBio 8:10.1128/mbio.01630-17.

Steele MI, Moran NA. 2021. Evolution of Interbacterial Antagonism in Bee Gut Microbiota Reflects Host and Symbiont Diversification. mSystems 6:10.1128/msystems.00063-21.

Steele MI, Motta EVS, Gattu T, Martinez D, Moran NA. 2021. The Gut Microbiota Protects Bees from Invasion by a Bacterial Pathogen. Microbiol. Spectr. 9:e00394–21.

Suria AM, Smith S, Speare L, Chen Y, Chien I, Clark EG, Krueger M, Warwick AM, Wilkins H, Septer AN. 2022. Prevalence and diversity of type VI secretion systems in a model beneficial symbiosis. Front. Microbiol. 13:988044.

Suzuki A, Hisamoto S, Sakamoto Y. 2025. Dynamics of the hindgut microbiota of the Japanese honey bees (*Apis cerana japonica*) throughout the overwintering period. PeerJ 13:e20050.

Thomas S, Holland IB, Schmitt L. 2014. The type 1 secretion pathway — the hemolysin system and beyond. Biochim. Biophys. Acta BBA - Mol. Cell Res. 1843:1629–1641.

Trunk T, Khalil HS, Leo JC. 2018. Bacterial autoaggregation. AIMS Microbiol. 4:140–164.

Ugarte-Ruiz M, Stabler RA, Domínguez L, Porrero MC, Wren BW, Dorrell N, Gundogdu O. 2015. Prevalence of type VI secretion system in spanish *Campylobacter jejuni* isolates. Zoonoses Public Health 62:497–500.

Unni R, Pintor KL, Diepold A, Unterweger D. 2022. Presence and absence of type VI secretion systems in bacteria. Microbiology 168:001151.

Vasimuddin Md, Misra S, Li H, Aluru S. 2019. Efficient architecture-aware acceleration of BWA-MEM for multicore systems. In: 2019 IEEE International Parallel and Distributed Processing Symposium (IPDPS). p. 314–324. Available from: https://ieeexplore.ieee.org/document/8820962

Veith PD, Glew MD, Gorasia DG, Reynolds EC. 2017. Type IX secretion: the generation of bacterial cell surface coatings involved in virulence, gliding motility and the degradation of complex biopolymers. Mol. Microbiol. 106:35–53.

Waksman G. 2025. Molecular basis of conjugation-mediated DNA transfer by gram-negative bacteria. Curr. Opin. Struct. Biol. 90:102978.

Wallner A, Moulin L, Busset N, Rimbault I, Béna G. 2021. Genetic diversity of type 3 secretion system in *Burkholderia* s.l. and links with plant host adaptation. Front. Microbiol. 12:761215.

Wang X, Sun B, Xu M, Qiu S, Xu D, Ran T, He J, Wang W. 2018. Crystal structure of the periplasmic domain of TssL, a key membrane component of type VI secretion system. Int. J. Biol. Macromol. 120:1474–1479.

Wangthaisong P, Piromyou P, Songwattana P, Wongdee J, Teamtaisong K, Tittabutr P, Boonkerd N, Teaumroong N. 2023. The type IV secretion system (T4SS) mediates symbiosis between *Bradyrhizobium sp.* SUTN9-2 and legumes. Appl. Environ. Microbiol. 89:e00040–23.

Weber BS, Ly PM, Irwin JN, Pukatzki S, Feldman MF. 2015. A multidrug resistance plasmid contains the molecular switch for type VI secretion in Acinetobacter baumannii. Proc. Natl. Acad. Sci. 112:9442–9447.

Wei Z, Zhang S, Wang X, Bai J, Wang H, Yang Y, Zhai J. 2025. Beyond survival to domination: *Brucella*’s multilayered strategies for evading host immune responses. Front. Microbiol. 16:1608617.

Whelan FJ, Rusilowicz M, McInerney JO. 2020. Coinfinder: detecting significant associations and dissociations in pangenomes. *Microb*. Genomics 6:e000338.

Whitfield GB, Brun YV. 2024. The type IVc pilus: just a Tad different. Curr. Opin. Microbiol. 79:102468.

Wong MJQ, Liang X, Smart M, Tang L, Moore R, Ingalls B, Dong TG. 2016. Microbial herd protection mediated by antagonistic interaction in polymicrobial communities. Appl. Environ. Microbiol. 82:6881–6888.

Wood DE, Lu J, Langmead B. 2019. Improved metagenomic analysis with Kraken 2. Genome Biol. 20:257.

Wu J, Lang H, Mu X, Zhang Z, Su Q, Hu X, Zheng H. 2021. Honey bee genetics shape the strain-level structure of gut microbiota in social transmission. Microbiome 9:225.

Xiong X, Wan W, Ding B, Cai M, Lu M, Liu W. 2024. Type VI secretion system drives bacterial diversity and functions in multispecies biofilms. Microbiol. Res. 279:127570.

Zboralski A, Biessy A, Filion M. 2022. Bridging the gap: type III secretion systems in plant-beneficial bacteria. Microorganisms 10:187.

Zhang J, Guan J, Wang M, Li G, Djordjevic M, Tai C, Wang H, Deng Z, Chen Z, Ou H-Y. 2022. SecReT6 update: a comprehensive resource of bacterial type VI secretion systems. Sci. CHINA Life Sci. 66:626–634.

Zhang L, Hinz AJ, Nadeau J-P, Mah T-F. 2011. *Pseudomonas aeruginosa* tssC1 links type VI secretion and biofilm-specific antibiotic resistance. J. Bacteriol. 193:5510–5513.

Zhang S, Wu F, Zhao H, Zhao L, Li D, Yang F, Liu L. 2025. Type IV secretion systems and conjugation in gram-negative pathogens. FASEB J. 39:e71116.

Zhang W, Zhang X, Su Q, Tang M, Zheng H, Zhou X. 2022. Genomic features underlying the evolutionary transitions of *Apibacter* to honey bee gut symbionts. Insect Sci. 29:259–275.

Zhang Z, Mu X, Cao Q, Shi Y, Hu X, Zheng H. 2022. Honeybee gut *Lactobacillus* modulates host learning and memory behaviors via regulating tryptophan metabolism. Nat. Commun. 13:2037.

Zheng H, Nishida A, Kwong WK, Koch H, Engel P, Steele MI, Moran NA. 2016. Metabolism of toxic sugars by strains of the bee gut symbiont *Gilliamella apicola*. mBio 7:10.1128/mbio.01326-16.

Zheng H, Perreau J, Powell JE, Han B, Zhang Z, Kwong WK, Tringe SG, Moran NA. 2019. Division of labor in honey bee gut microbiota for plant polysaccharide digestion. Proc. Natl. Acad. Sci. 116:25909–25916.

Zheng H, Powell JE, Steele MI, Dietrich C, Moran NA. 2017. Honeybee gut microbiota promotes host weight gain via bacterial metabolism and hormonal signaling. Proc. Natl. Acad. Sci. 114:4775–4780.

Zhou N, Zheng Q, Liu Y, Huang Z, Feng Y, Chen Y, Hu F, Zheng H. 2025. Strain diversity and host specificity of the gut symbiont *Gilliamella* in *Apis mellifera, Apis cerana* and *Bombus terrestris*. Microbiol. Res. 293:128048.

Zoued A, Brunet YR, Durand E, Aschtgen M-S, Logger L, Douzi B, Journet L, Cambillau C, Cascales E. 2014. Architecture and assembly of the Type VI secretion system. Biochim. Biophys. Acta BBA - Mol. Cell Res. 1843:1664–1673.

