## Supplementary material for "Vertical inheritance and loss-driven evolution of secretion systems in the bee gut microbiota": Fig. S2

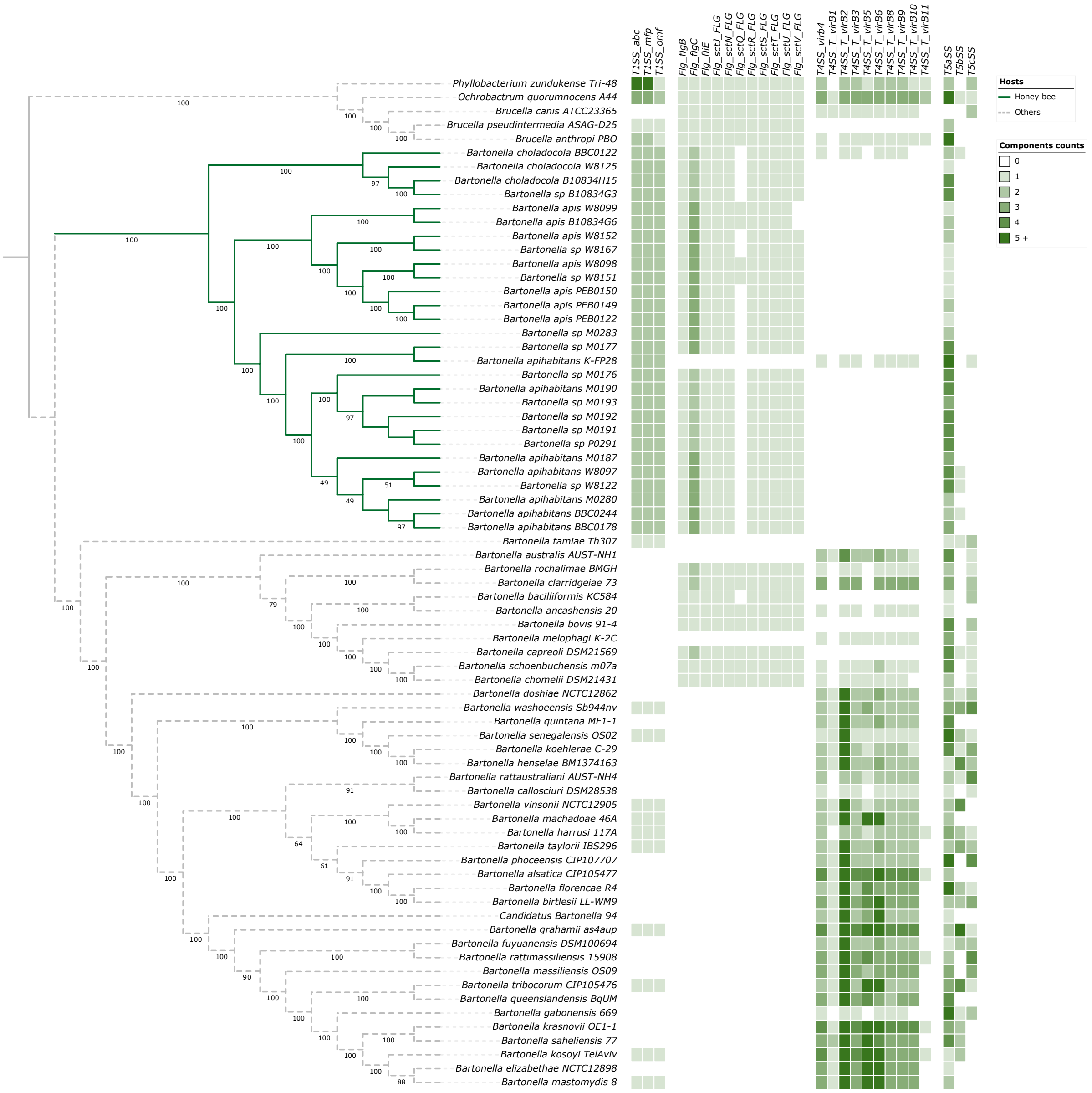


**Figure S2. Detected secretion system components of the bee microbiota lineage *Bartonella* (solid branches) and related bacteria of the *Bartonellaceae* family.** Cladogram depicts strain relationships as inferred from a maximum likelihood phylogenomic tree generated using 247 single-copy orthologous genes under the GTR+F+I+I+R7 model with 1000 non-parametric bootstrap replicates, shown as percentages at nodes.
