## Supplementary material for "Vertical inheritance and loss-driven evolution of secretion systems in the bee gut microbiota": Fig. S4

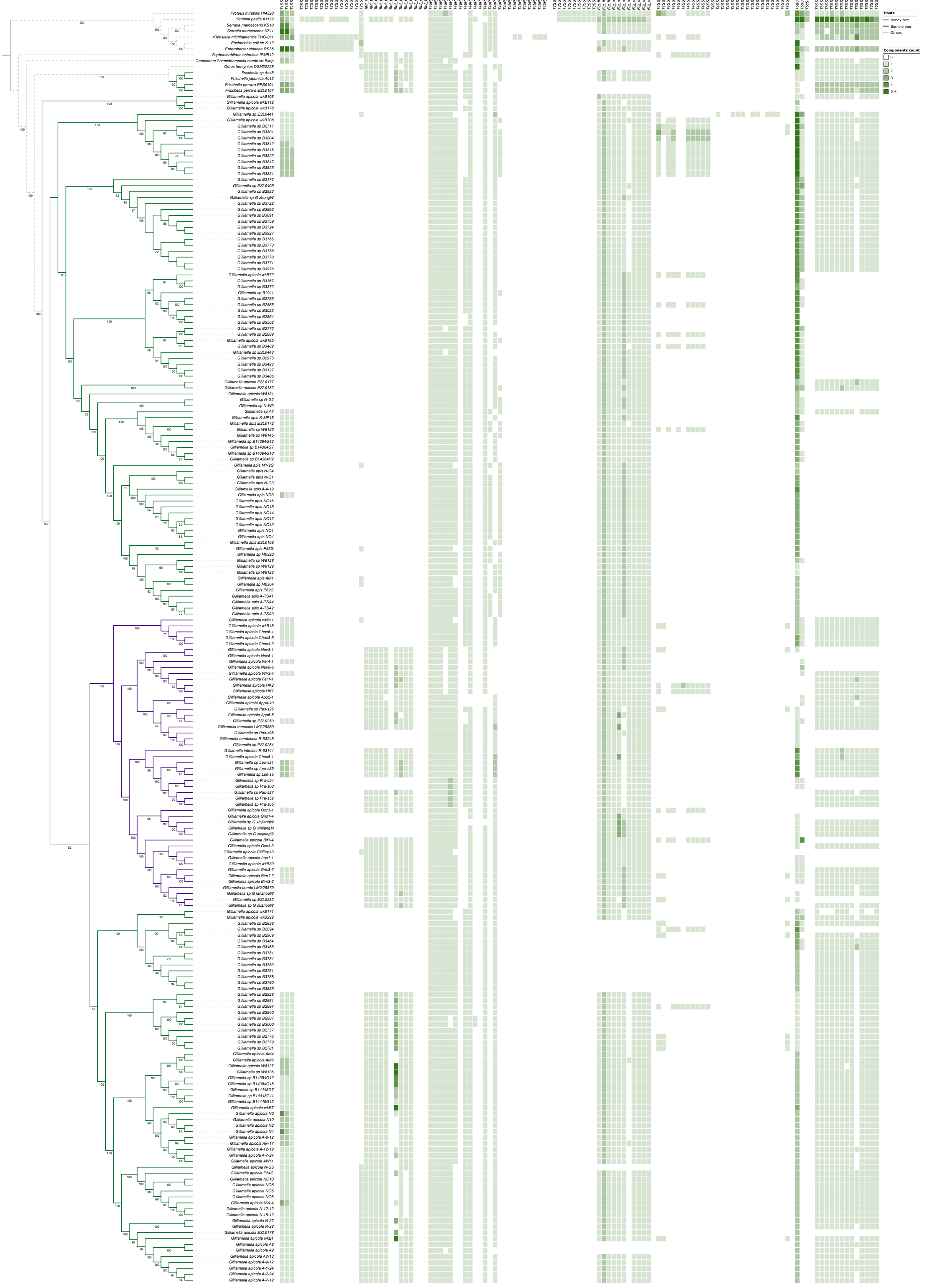

**Figure S4. Detected secretion system components of the bee microbiota lineages *Frischella* and *Gilliamella* (solid branches) and related bacteria of the *Orbaceae* family.** Cladogram depicts strain relationships as inferred from a maximum likelihood phylogenomic tree generated using 337 single-copy orthologous genes under the GTR+F+I+I+R9 model with 1000 non-parametric bootstrap replicates, shown as percentages at nodes.
