## Supplementary material for "Vertical inheritance and loss-driven evolution of secretion systems in the bee gut microbiota": Fig. S6

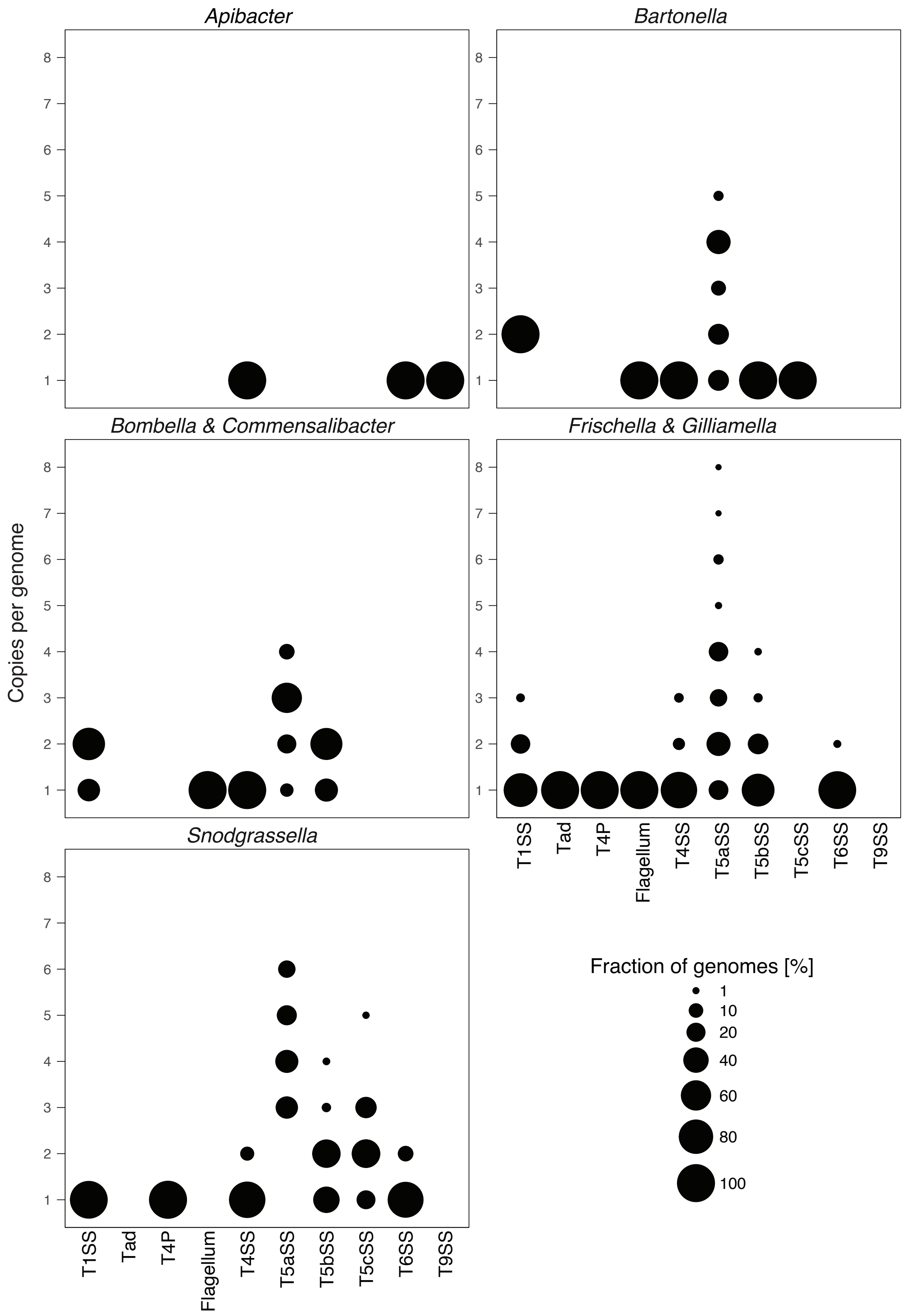


**Figure S6. Distribution of the number of each secretion system detected in the genomes across each bee-associated bacterial lineage.** Bubble size reflects the proportion (%) of bacteria that encode the indicated count for each system.
