## Supplementary material for "Vertical inheritance and loss-driven evolution of secretion systems in the bee gut microbiota": Fig. S8

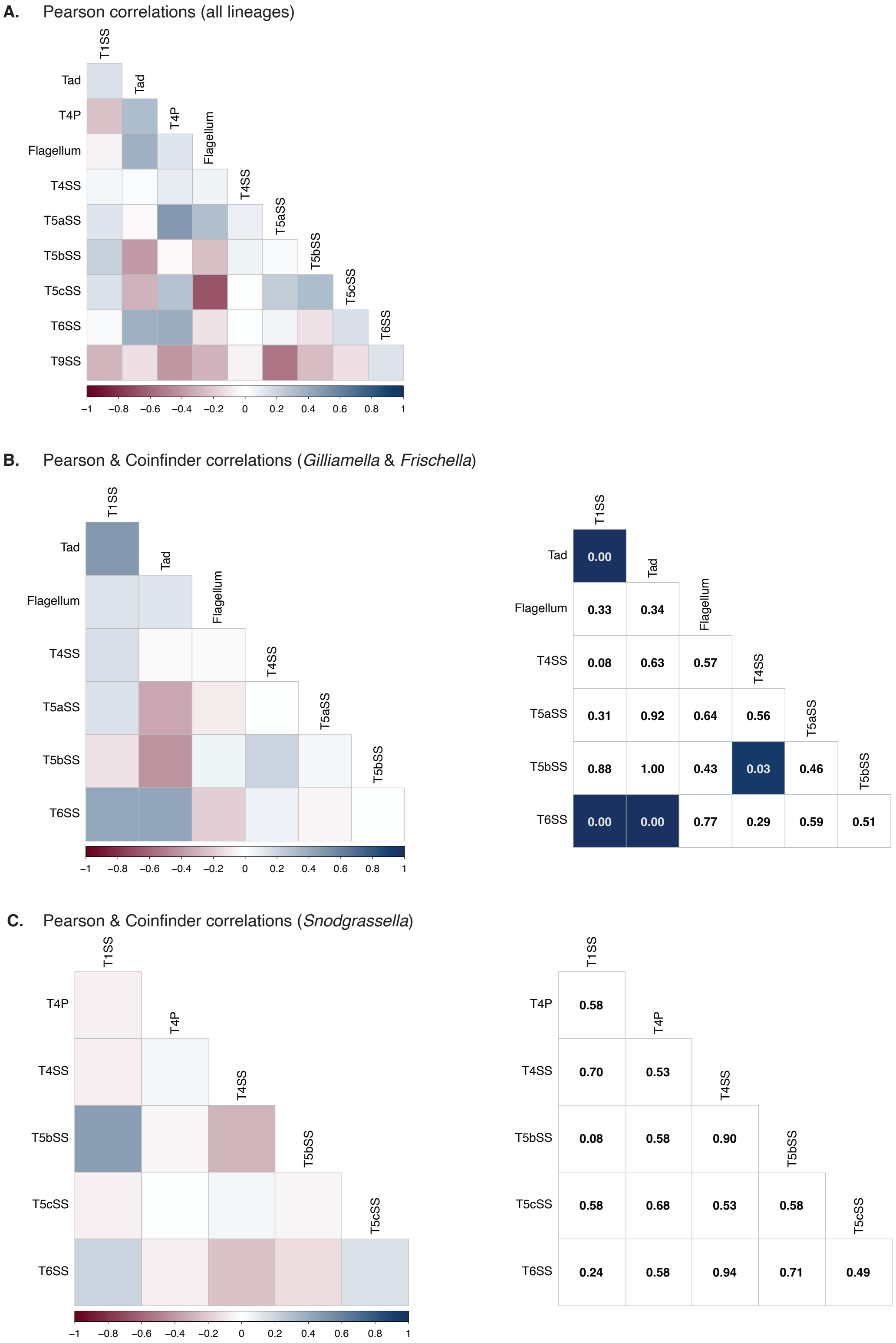


**Figure S8. Correlation of secretion system presence/absence across bee gut bacterial genomes. (A)** Correlation analysis across all genomes. The presence and absence of each secretion system and appendage were encoded as binary variables (1 = present, 0 = absent), and systems lacking variation were excluded. Pairwise associations were calculated using Pearson's correlation coefficient, and the resulting correlation matrix is shown as a heatmap. Colour intensity indicates the strength and direction of the correlation, with positive values representing systems that tend to co-occur and negative values indicating mutually exclusive patterns. **(B)** Correlation (left side matrix) and phylogeny-aware co-occurrence (*p*-value ≤ 0.05, coloured squares, right side matrix) in *Gilliamella* & *Frischella*, and **(C)** *Snodgrassella* genomes.
