## Supplementary material for "Vertical inheritance and loss-driven evolution of secretion systems in the bee gut microbiota": Fig. S9

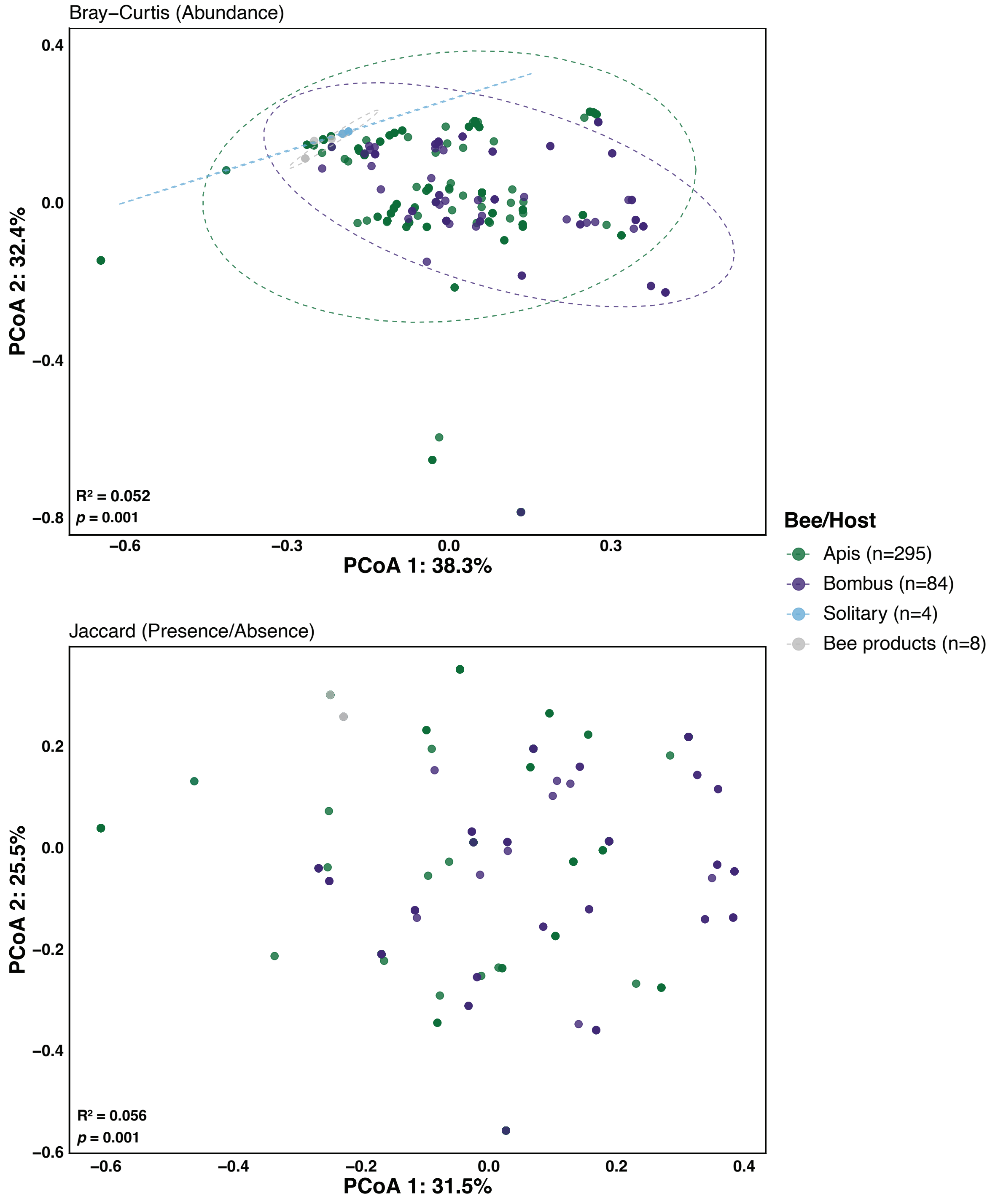


**Figure S9. Secretion system diversity across bee hosts.** Beta diversity was assessed using Bray–Curtis dissimilarity based on Hellinger-transformed secretion system abundances and Jaccard distance based on presence/absence data. Ordinations visualized using principal coordinates analysis (PCoA), with the percentage of variance explained by each axis indicated. Differences in secretion system composition among lineages were tested using PERMANOVA (999 permutations), with effect sizes reported as $R^2$. Ellipses represent 95% confidence intervals around host centroids.
