## Supplementary material for "Vertical inheritance and loss-driven evolution of secretion systems in the bee gut microbiota": Fig. S10

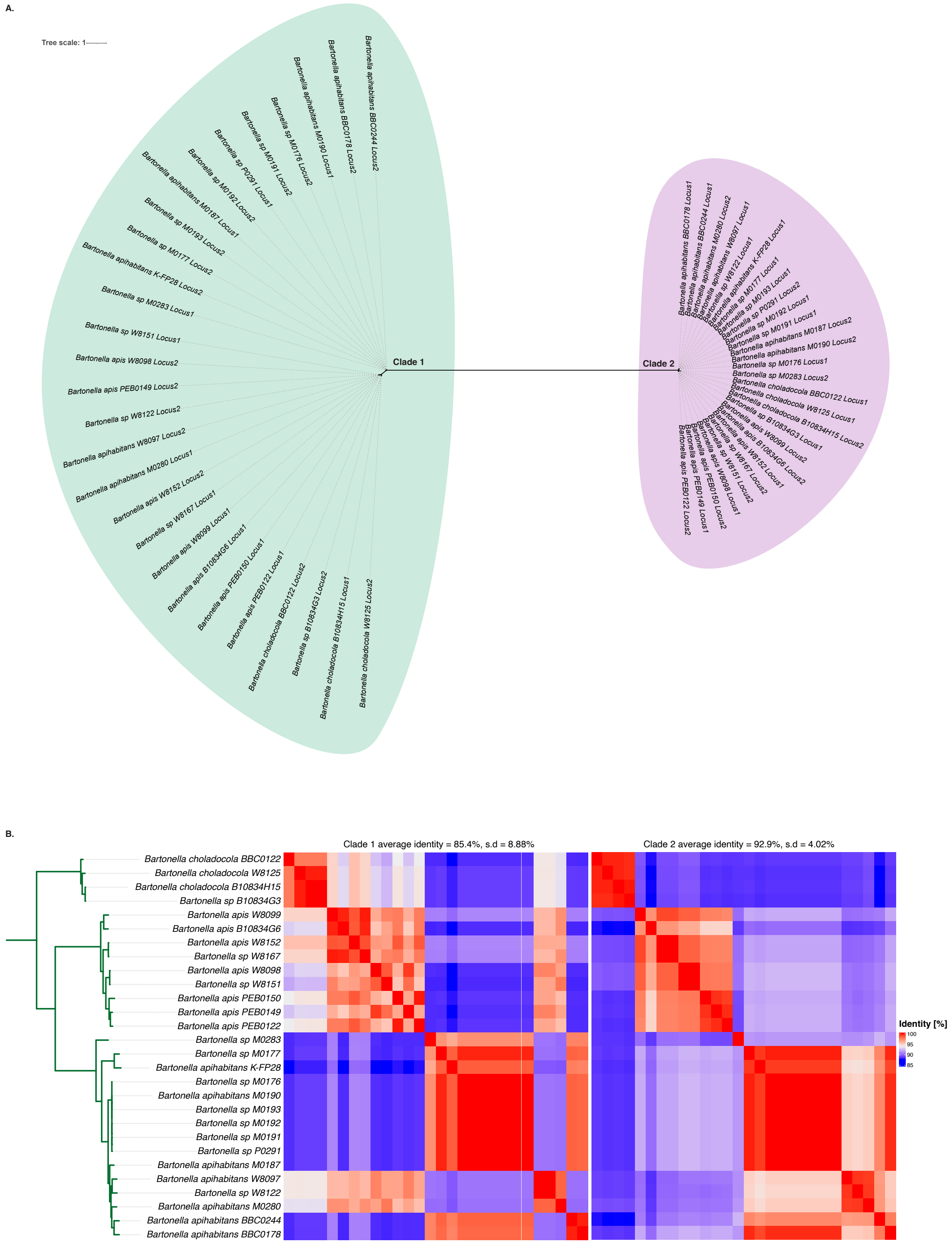


**Figure S10. The type I secretion systems of the bee-associated *Bartonella* lineage. (A)** Maximum likelihood phylogeny of concatenated *abc*, *mfp* and *omf* genes of the T1SS, showing the two distinct copies of the system, labelled clade 1 and clade 2. Tree was generated under the LG+F+I+G4 model with 1000 non-parametric bootstrap replicates. **(B)** Pairwise amino acid identity matrix of the T1SS within the two clades. Strains (left labels) are ordered by their lineage phylogeny (Fig. S2), and strain identity of the columns (left to right) are in the same order as the row labels (top to bottom).
