## Supplementary material for "Vertical inheritance and loss-driven evolution of secretion systems in the bee gut microbiota": Fig. S11

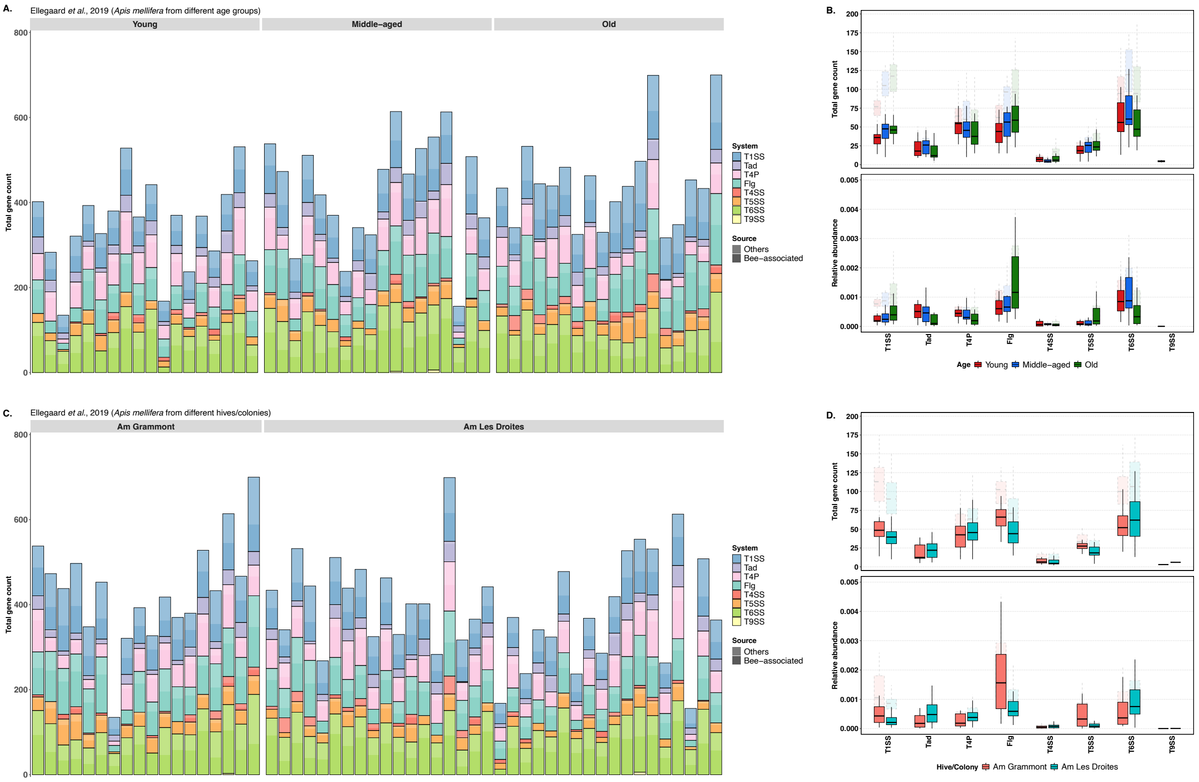


**Figure S11. Distribution of secretion system and appendages gene counts in *Apis mellifera* gut bacterial metagenomes from Ellegaard *et al.* (2019).** Stacked bar plot showing total gene counts per secretion system in individual metagenomes, grouped by **(A)** bee age and **(C)** hive/colony. Each bar represents one metagenome, coloured by system, with darker shading indicating the fraction attributed to bee-associated taxa. Boxplots showing total gene counts (top) and relative abundance (bottom) of each secretion system across **(B)** age groups and **(D)** bee hives/colonies. Dashed pale outlines represent all detected genes, and solid filled boxes represent only genes from bee-associated bacteria.
