## Supplementary material for "Vertical inheritance and loss-driven evolution of secretion systems in the bee gut microbiota": Fig. S13

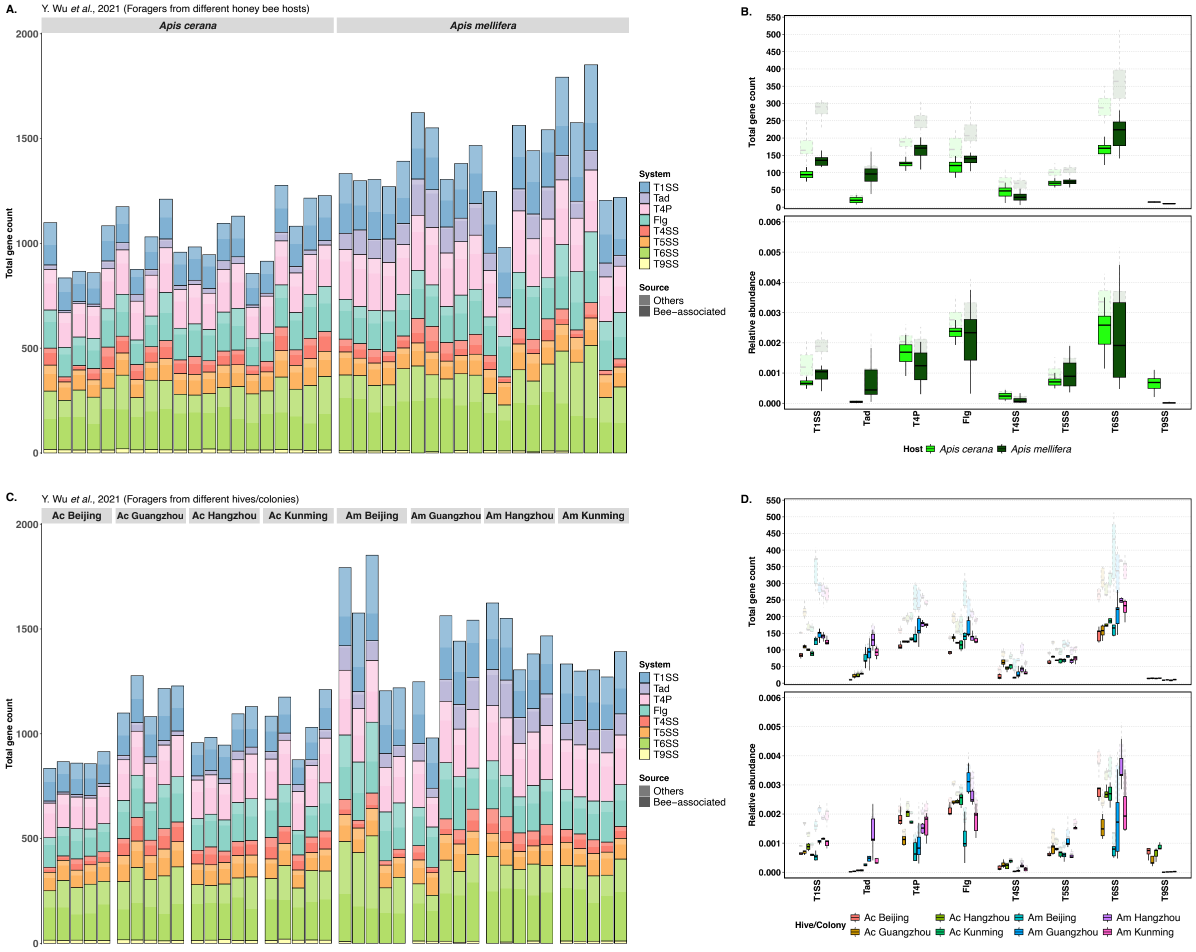


**Figure S13. Distribution of secretion system and appendages gene counts in forager (old) *Apis mellifera* and *Apis cerana* gut bacterial metagenomes from Y Wu *et al.* (2021).** Stacked bar plot showing total gene counts per secretion system in individual metagenomes, grouped by **(A)** host species and **(C)** bee hive/colony. Each bar represents one metagenome, coloured by system, with darker shading indicating the fraction attributed to bee-associated taxa. Boxplots showing total gene counts (top) and relative abundance (bottom) of each secretion system across **(B)** host species and **(D)** bee hives/colonies. Dashed pale outlines represent all detected genes, and solid filled boxes represent only genes from bee-associated bacteria.
