## Supplementary material for "Vertical inheritance and loss-driven evolution of secretion systems in the bee gut microbiota": Fig. S14

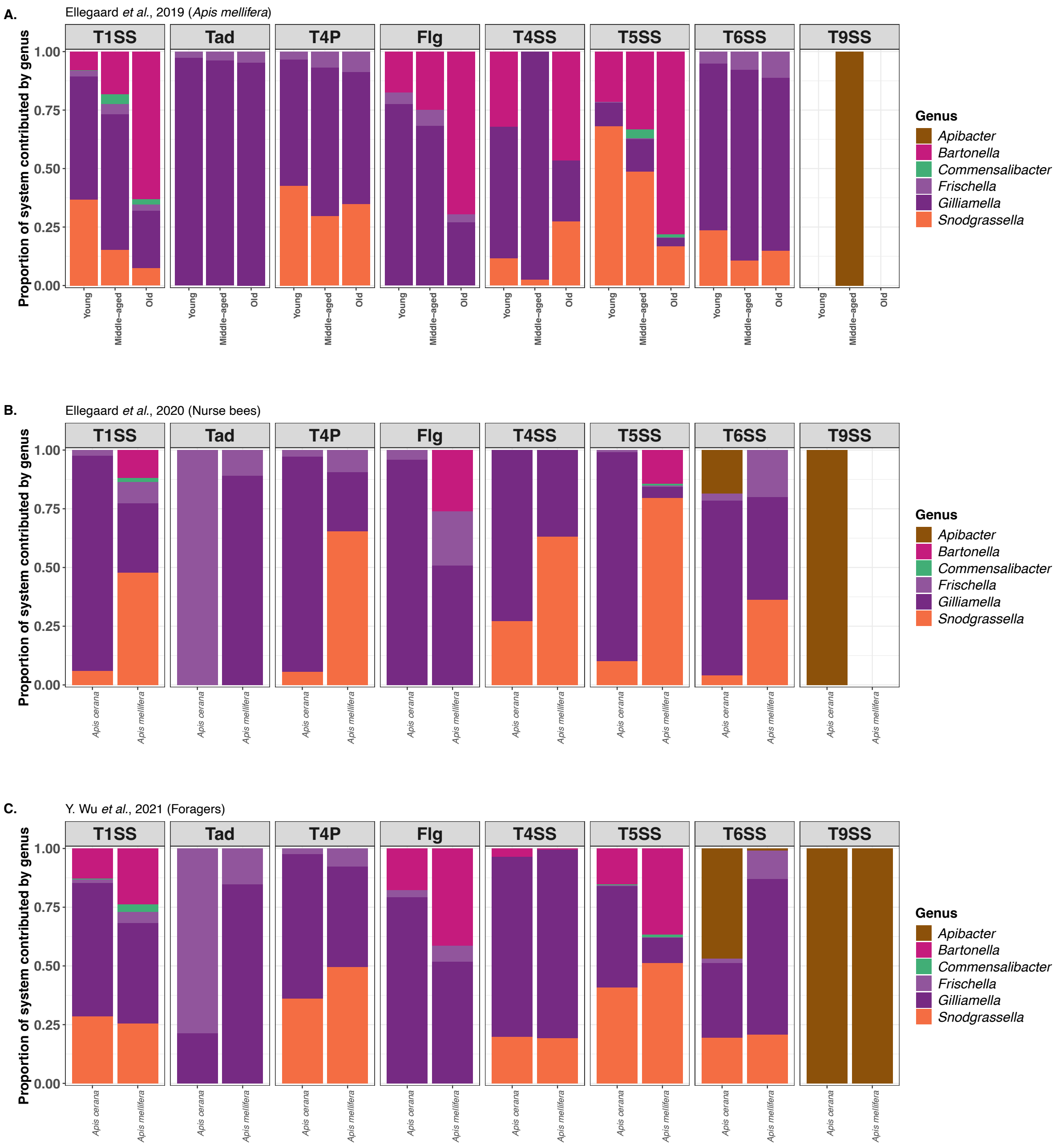


**Figure S14. Taxonomic profiling of secretion system–associated contigs across honey bee metagenomes.** Metagenomic reads were mapped to contigs corresponding to each secretion system across datasets obtained from **(A)** young (10 days old), middle-aged (22-24 days) and old (≥48 days) *Apis mellifera*; **(B)** from nurses (young) *Apis cerana* and *A. mellifera*; and **(C)** from foraging (old) *A. cerana* and *A. mellifera*.
