## Supplementary material for "Vertical inheritance and loss-driven evolution of secretion systems in the bee gut microbiota": Fig. S15

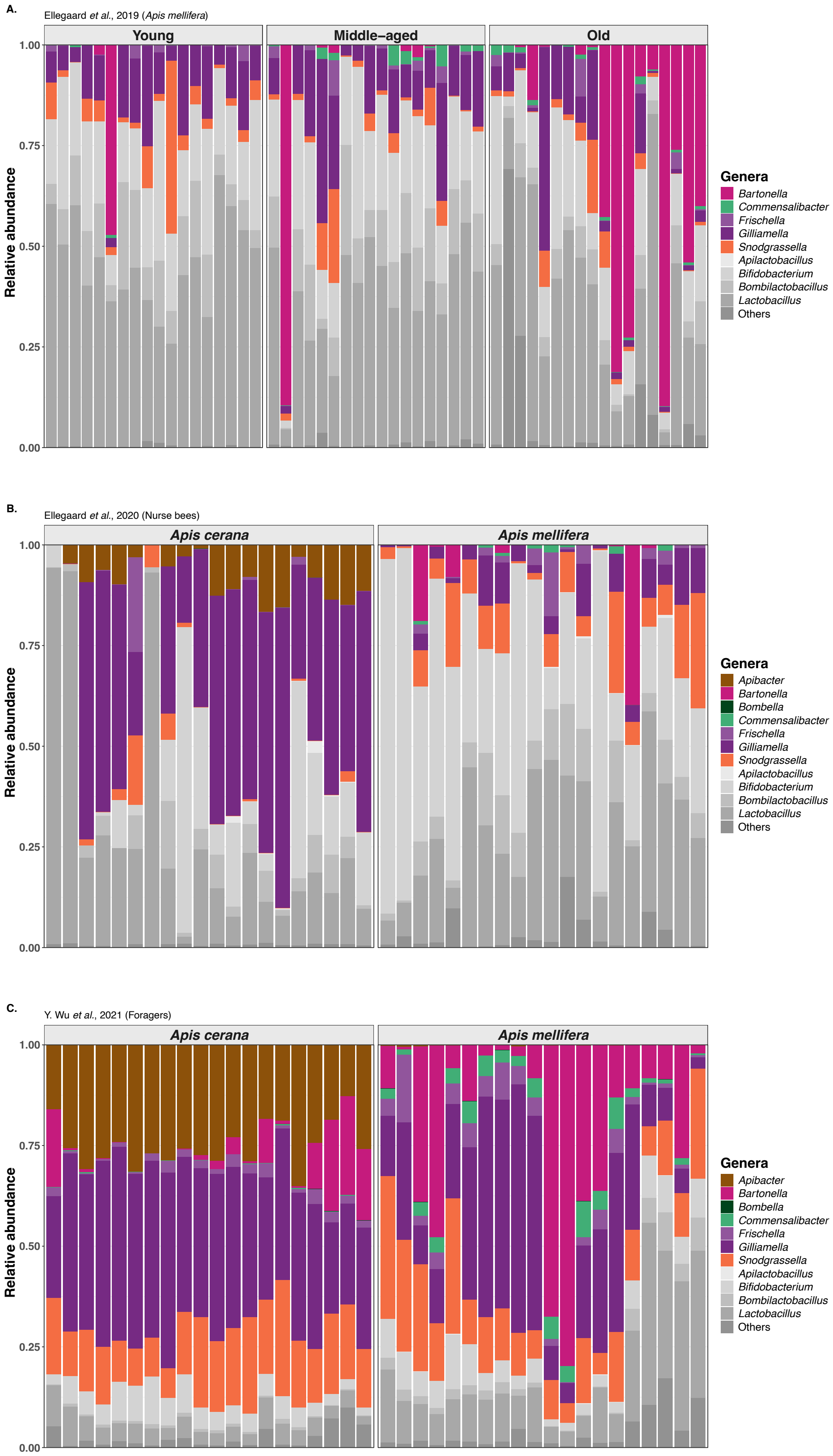


**Figure S15. Taxonomic composition of bee gut metagenomes across the three studies**. Gram-negative core bee bacterial genera highlighted/coloured and Gram-positive/other bacteria shown in shades of grey. Relative abundance of bacterial genera in **(A)** *Apis mellifera* of different ages (Ellegaard *et al.*, 2019), **(B)** nurse (young) *Apis cerana* and *A. mellifera* bees (Ellegaard *et al.*, 2020), and **(C)** foraging (old) *A. cerana* and *A. mellifera* bees (Wu *et al.*, 2021).
