## Supplementary material for "Vertical inheritance and loss-driven evolution of secretion systems in the bee gut microbiota": Fig. S16

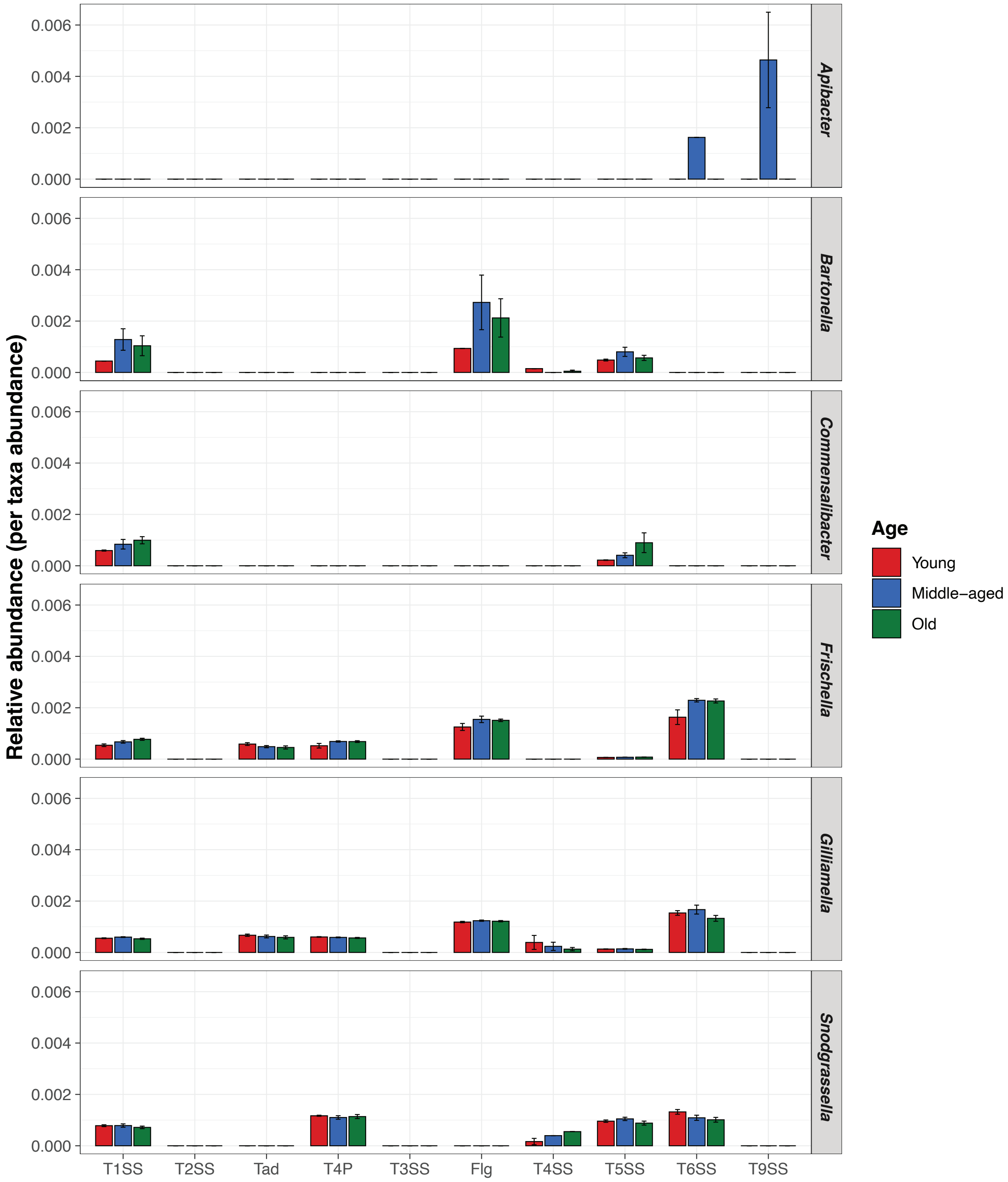


**Figure S16. Relative abundance of secretion systems across bee gut bacterial genera in *Apis mellifera* of different ages (Ellegaard *et al.*, 2019).** Bar plots show the mean relative abundance of secretion systems normalized by the corresponding genus abundance for each age category of bees, with error bars representing the standard error of the mean.
